# A novel method for generating 3D constructs with branched vascular networks using multi-materials bioprinting and direct surgical anastomosis

**DOI:** 10.1101/2021.03.21.436268

**Authors:** Xin Liu, Xinhuan Wang, Liming Zhang, Lulu Sun, Heran Wang, Hao Zhao, Zhengtao Zhang, Yiming Huang, Jingjinqiu Zhang, Biaobiao Song, Chun Li, Hui Zhang, Song Li, Shu Wang, Xiongfei Zheng, Qi Gu

**Affiliations:** State Key Laboratory of Membrane Biology, Institute of Zoology, Chinese Academy of Sciences, Beijing 100101, P. R. China; Savaid Medical School, University of Chinese Academy of Sciences, Beijing 100049, P. R. China; Shenyang Institute of Automation, Chinese Academy of Sciences, Shenyang 110169, P. R. China; Institute of Chemistry, Chinese Academy of Sciences, Beijing, 100190 P. R. China; State Key Laboratory of Cell Biology, CAS Center for Excellence in Molecular Cell Science, Shanghai Institute of Biochemistry and Cell Biology, Chinese Academy of Sciences, Shanghai 200031, P. R. China; University of Science and Technology of China, Hefei, 230026, P. R. China

**Keywords:** 3D printing, multivascular network, low viscosity, GelMA, hUVEC, 3D bioprinting, vasculature, vascularization, perfusion, transplantation

## Abstract

Vessels pervade almost all body tissues, and significantly influence the pathophysiology of human body. Previous attempts to establish multi-scale vascular connection and function in 3D model tissues using bioprinting have had limited success due to the incoordination between cell-laden materials and stability of the perfusion channel. Here, we report a methodology to fabricate centimetre-scale vascularized soft tissue with high viability and accuracy using multi-materials bioprinting involving inks with low viscosity and a customized multistage-temperature-control printer. The tissue formed was perfused with branched vasculature with well-formed 3D capillary network and lumen, which would potentially supply the cellular components with sufficient nutrients in the matrix. Furthermore, the same methodology was applied for generating liver-like tissue with the objective to fabricate and mimic a mature and functional liver tissue, with increased functionality in terms of synthesis of liver specific proteins after *in vitro* perfusion and *in vivo* subperitoneal transplantation in mice. Moreover, to establish immediate blood perfusion, an elastic layer was printed wrapping sacrificial ink to support the direct surgical anastomosis of the carotid artery to the jugular vein. Our findings highlight the support extended by vasculature network in soft hydrogels which helps to sustain the thick and dense cellularization in engineered tissues.

## 1. Introduction

Three-dimensional (3D) bioprinting techniques have significantly facilitated the process of fabrication of complex, heterocellular soft artificial tissues *in vitro*, which combine polymeric biomaterials and cells.^[1–3]^ During the process of bioprinting, the bioinks provide protection to the cellular component, ensuring high cell viability, while also mimicking the extracellular matrix to promote bioactivity.^[4–6]^ Although bioprinting technology has shown great potential in tissue engineering, the thickness of constructed tissues was limited to several hundred micrometers due to restricted oxygen and nutrient diffusion, which is integral in maintaining cell viability and proliferation.^[7–8]^ In highly vascularized tissues, such as liver and kidney, the formation of new blood vessels is essential for growth beyond the diffusion limit.^[9–10]^ Therefore, building multi-branched perfusable vascular networks is critical to the fabrication of thick tissue constructs.

Recent advances in 3D tissue fabrication have led to efficient bioprinting of blood vessels.^[11–12]^ The strategies followed in these studies can be classified into two main groups: (i) scaffold-based approach, and (ii) scaffold-free bioprinting of vascular constructs.^[13]^ The first approach can be divided into three major bioprinting modalities which include extrusion-based bioprinting, droplet-based bioprinting, and laser-based bioprinting.^[14–15]^ Extrusion-based bioprinting enables fabrication of macro-vascular constructs (in the order of magnitude of a few centimeters), which allows printing with fugitive inks with subsequent remove for achieving a distinct vascular pattern. For demonstrating this method, water-soluble sugar ink was first utilized.^[16]^ The capillaries can be fabricated with diameters as small as 150 m. Recently, an interconnected vascular network that perfuses a centimeter-scale engineered osteogenic construct was generated.^[17]^ The fugitive ink Pluronic F127 was dissolved, and endothelial cells (ECs) were used to create a vascular bed by seeding the channels. Alginate and gelatin have also been used as fugitive inks in recent studies.^[18–20]^ However, cell-laden inks have the limitations of low cell viability (due to shear damage from ink extrusion) and exhibit poor resolution of the final vascular pattern formed (hundreds of micrometer voxels).^[21–22]^ For scaffold-free bioprinting, cells encapsulated in hydrogels or decellularized matrix components could be used to fabricate micro-vascular network, which otherwise relies on ECs to form new vessel by physiological mechanisms.^[23]^ Recently, pre-vascularized spheroidal assemblies with elaborate branching have been used to form vascular networks.^[24]^ This proves that successful cellular assembly can be achieved after bioprinting by accelerated vascularization, which leads to the maturation of the tissue constructs.^[14, 25]^ Furthermore, this method can mimic tissue regeneration and development; however, the vessels formed are limited in size and do not exceed the micrometer scale.^[25]^

The bioprinted tissue constructs with blood microvessels would eventually be implanted, and therefore they should be suitable for surgical anastomosis to the host vasculature after implantation.^[26]^ Synthetic biodegradable microvessel microfluidic scaffolds provide sufficient structural support and have been successfully integrated with the host vasculature.^[27–28]^ However, soft hydrogel with vascular networks lacked sufficient mechanical properties, which severely limited their utilization as a load-bearing construct in tubular tissue regeneration.^[16]^ So far, no research effort has been devoted to integration and implantation of hydrogel-based hollow vascular network into host vasculature. Some efforts have used sacrificial laser-sintered for constructing vascular networks, but this indirect printing method was detrimental to integrated bioprinting using multi-materials.^[29]^ Therefore, we propose a method for bioprinting vasculature structure using elastic hydrogel and cell-laden hydrogel for enhanced mechanical support and biological activity. Cell-laden hydrogels may potentially decrease the damage during bioprinting and solidify in a cell-friendly environment, which may result in optimal cell proliferation and tissue remodeling.

In this study, we report the use of 3% GelMA with fibrin, as the cell-laden biomaterial for extrusion and bioprinting of vascularized tissue. With gelatin as fugitive inks, we printed HepG2 aggregates (HAs) with human umbilical vein endothelial cells (HUVECs) and mesenchymal stem cells (MSCs) as mixture to fabricate a vascularized hepatorganoid tissue (HOs), with proper endothelialization. Moreover, fabrication of functional 3D printed liver tissue with normal hepatocytic function was explored in this study, which involved probing liver-specific gene expression and albumin secretion *in vitro*, and *in vivo* subperitoneal transplanted in mice (**Figure 1A**). Furthermore, we leveraged this methodology combining 5% GelMA as inner elastic inks and external elastic inks to establish surgical anastomosis of the tissue, which has been rarely reported previously (Figure 1B).

**Figure 1.**
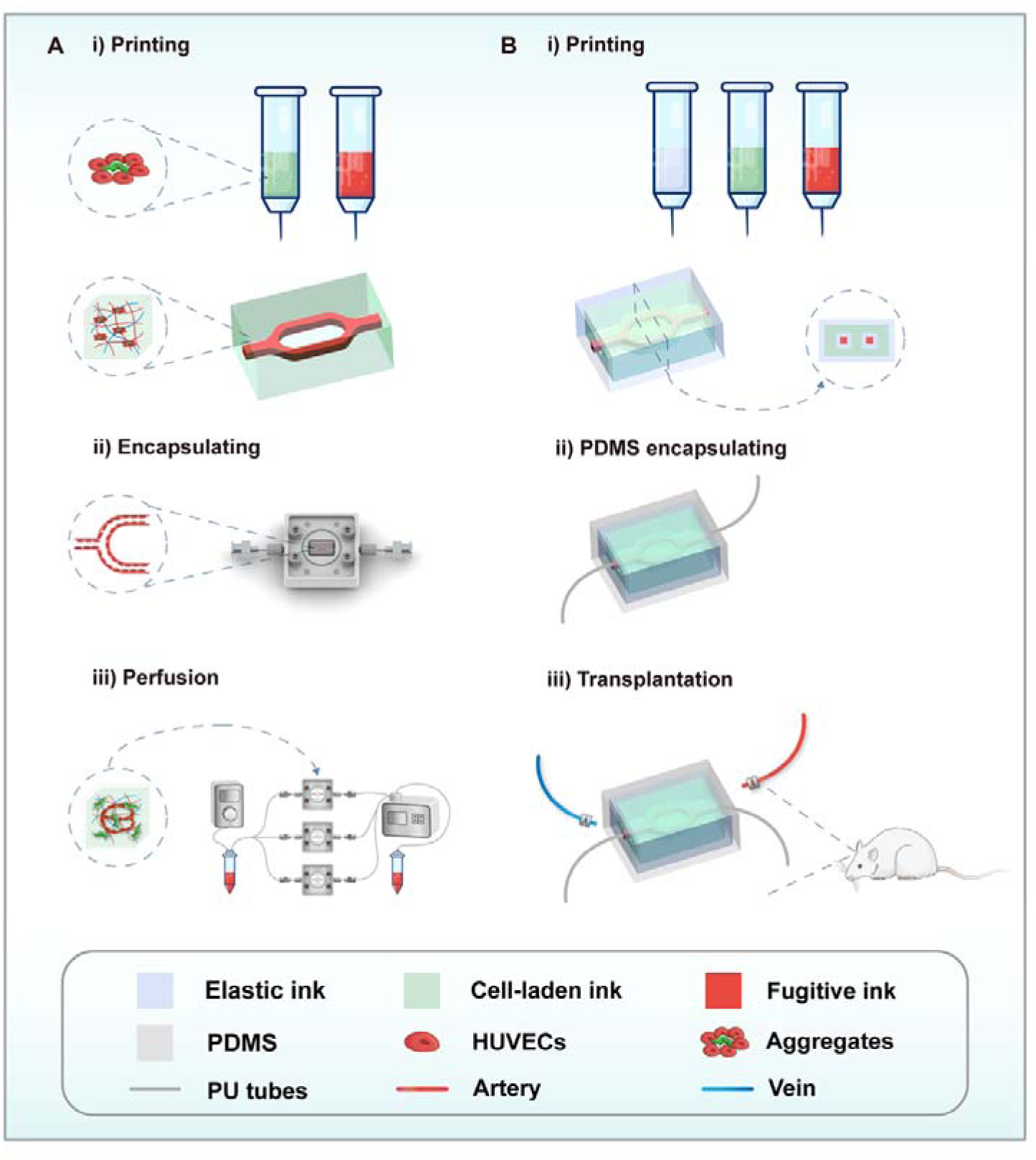
Schematic representation of multi-materials bioprinting strategies used in this study. A) Multi-materials bioprinting was performed using cell-laden inks and fugitive inks to fabricate 3D hepatic tissue with proper vascularization, and hepatocytic function *in vitro*. B) Multi-materials bioprinting was performed using cell-laden inks, fugitive inks, and elastic inks to fabricate tissue and optimal perfusion was achieved *in vivo* by direct surgical anastomosis to host vasculature (artery to vein).

## 2. Results and Discussion

### 2.1. 3D printing of vascular constructs

The viscosity of bioink is one of the important factors directly influencing the process of printing. When the viscosity of bioink is less (i.e.<5 Pa·s), the precision of printing is low. However, when the viscosity is high, it cannot protect cells from high shear stress of printing, which results in low cell survival rates.^[30]^ Recent studies have demonstrated the suitability of GelMA hydrogels with low concentration (<5% w/v) to perform as cell-based bioinks due to their high cell stability and viability.^[31–32]^ In this study, we evaluated the viscosity of GelMA under different temperature conditions (**Figure 2A**). When the GelMA concentration was 1% or 2%, its viscosity was close to zero and its properties were hardly affected by temperature. However, when the concentration was 3%, 4%, or 5%, its viscosity increased with the increase in GelMA concentration (at 5°C), which gradually decreased with the increase in temperature. However, when the temperature exceeded 25°C, the viscosity of GelMA approached zero. Moreover, the addition of 0.25% fibrin had no effect on the viscosity of GelMA. In order to visualize the change in viscosity of different concentrations of GelMA under different temperature conditions, the viscosity data were fitted to establish a three-dimensional heat map representing viscosity-temperature-concentration (Figure 2B). The red area represents the lower viscosity, and the blue area represents higher viscosity. For printability, 3% is the lowest concentration of GelMA with controllable viscosity at temperatures above zero.

**Figure 2.**
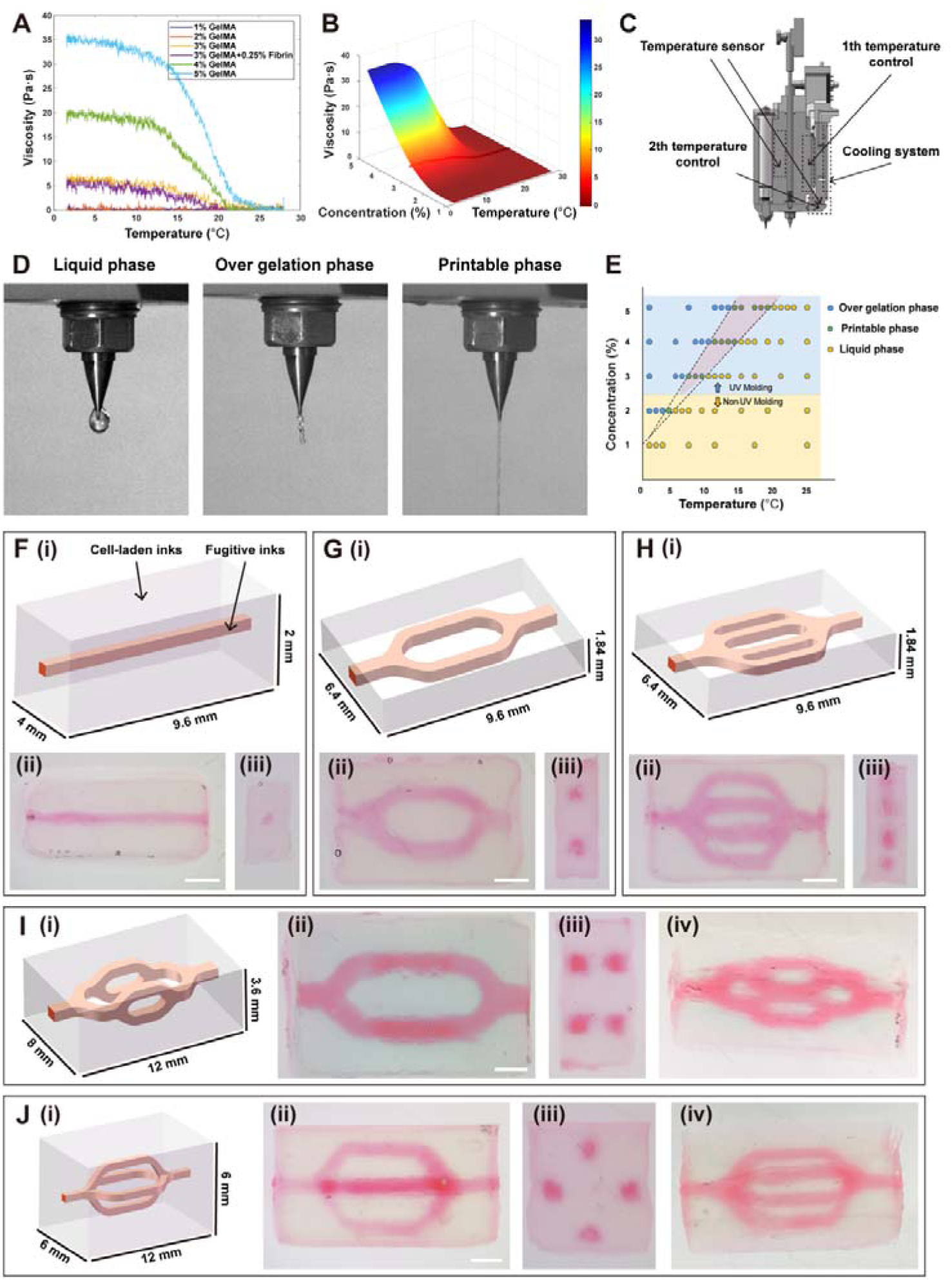
A) Viscosity test indicating the viscosity of GelMA and GelMA-fibrin at different concentrations. B) Three-dimensional graph indicating viscosity-temperature of GelMA and GelMA-fibrin at different concentrations. C) Visual representation of the printing system. D) Extrusion state of GelMA at different concentrations and temperatures, showing liquid phase, over gelation phase, and printable phase at different temperatures. E) Graphical representation of the extrusion state of GelMA at different concentrations and temperatures showing the printable region. F-J) Schematic illustrations and optical images of different constructs printed using transparent ink along with cell-laden ink, red ink along with fugitive ink, respectively. (i) schematic illustrations, (ii) top view, (iii) side view, (iv) stereogram. Scale bar represents 1 mm.

GelMA is highly sensitive to temperature and light.^[33]^ In order to achieve high precision, it is necessary to use GelMA in its printable phase and UV photocrosslinking was used after completion of the printing. The printable phase and UV crosslinking are two basic requirements for GelMA-based printing. The extrusion phase of GelMA recorded and the printing head (BiopHead) used in this study are shown in Figure 2C and Figure 2D. The gel point temperature T_0_ was measured from rheological tests (whereby storage modulus was equal to loss modulus). By setting the primary temperature of the printing head to be slightly higher than the gel point temperature T_0_, the liquid phase of GelMA was maintained. The extrusion phase of GelMA was tested by adjusting the secondary temperature of the BiopHead. The extrusion phase of GelMA was analyzed through a high-speed camera (Figure 2D). GelMA remained in over gelation phase when the temperature was above T_0_, with the extruded GelMA being distorted and irregular. GelMA remained in liquid phase when the temperature was below T_0_, extruding the GelMA as a droplet at the tip of the needle. When the temperature condition was set correctly, the GelMA was in printable phase with the extruded GelMA being smooth with superior printing performance. As shown in Figure 2C and Figure 2E, the printability of low concentration GelMA is one of the goals of the experiment, and it is necessary to explore the printability of low concentration GelMA. Due to the temperature and light sensitive properties of GelMA, it is need to be cooled in the printhead to achieve pre-gel extrusion and molding, and the physical gelatinized construction is chemically crosslinked by UV lamps and cultures at 37°C to maintain shape accuracy. Therefore, the maintenance of pre-gel state and purple diplomatic post-union structure are two basic requirements for GelMA. First, the extrusion status of the GelMA was recorded. The printhead used in the test was shown in Figure 2D. The gel point temperature T_0_ (the storage modulus is equal to loss modulus) was measured based on the rheological properties of bioink. The liquid state of the bioink in the print silo was maintained by setting the first stage temperature of the printhead slightly higher than the gel point temperature T_0_. The extrusion state of the bioink was tested by adjusting the secondary temperature of the printhead. The green circle in the figure represents the pre-gel state of the bioink. Secondly, the maintenance effect of the construction after the purple couplet was tested. When the concentration of GelMA is lower than 2%, the printing requirements in the state of physical gel could be achieved by lowering the temperature. However, the structure of the chemical crosslink will melt and collapse in the environment of 37□, which could not meet the stability requirements. The solidification and forming effect of bioink in the blue area of the test diagram is better. The above experimental results show that the concentration and temperature parameters of bioink in the pink range of the map are printable regions.

Figure 2C and Figure S1 show the BiopHead printing system used in the experiment, which includes air-driven silo, electric-driven silo, primary temperature control device, secondary temperature control device, and a cooling system. BiopHead can achieve a temperature gradient through a two-stage temperature control system, which allows the bioink in the printing silo to be in a liquid phase, and remain in a printable phase in the printing needle. BiopHead can run in two extrusion modes, namely the electric extrusion mode and the pneumatic extrusion mode. Electric extrusion is based on volume change. In this mode, direct-current motor drives the screw structure to rotate, which in turn drives the push rod (the pushing device behind the syringe) to move linearly. The underlying reason for the high precision in the electric extrusion mode is the precision of the screw displacement. It is difficult to achieve high-precision control of the print volume when the viscosity of the bioink is low, i.e., the bioink is in a thin state. Also, since the print silo of the electric extrusion is smaller, the displacement of the push rod has less influence on the extrusion volume. The pneumatic extrusion is based on pressure change. The pressure of the rear end of the silo is changed by an air compressor (cylinder) which causes the extrusion of the bioink. In the pneumatic extrusion mode, the volume extruded is larger, which makes it ideal for large-volume printing. Controlling the pressure of the ink is difficult, hence, printing accuracy is not guaranteed. Therefore, electric extrusion is used for high-precision printing, while pneumatic extrusion is used for large-scale printing involving larger volumes (e.g.<5 ml). In this study, GelMA was printed using an electric drive, and the temperature control system was adjusted to maintain the bioink in the gel state. The temperature of the printing silo was controlled by the primary temperature control system to keep bioink in the printing silo in liquid phase. The temperature of the printing needle was controlled by the secondary temperature control system to maintain the bioink in a printable phase suitable for printing. The extrusion state of the bioink was captured by a high-speed camera. When the temperature was too low, the bioink was in over gelation phase. When the temperature was too high, the bioink was in a liquid phase, and when the temperature conditions were right, the bioink was in a printable phase.

To engineer and fabricate vascularized tissue constructs, several channel configuration structures were designed. First, one straight channel within a matrix was designed, which was printed with both cell-laden inks and sacrificial inks Figure 2F. The cell-laden inks and fugitive inks were printed on a glass substrate with dimensions of 9.6 mm × 4 mm × 2 mm. Next, photocrosslinking was achieved by exposing the printed constructs to UV light for 2 mins. Then, the fugitive ink was removed from the thick tissue by heating to ∼37°C, whereby it undergoes a gel-to-fluid transition. Furthermore, we printed several different constructs to demonstrate the formation of stable vascularized tissues, including a one-to-two-channeled structure Figure 2G, a one-to-four-channeled structure Figure 2H, a one-to-two-to-four-channeled 3D structure Figure 2I, and a one-to-four-channeled 3D structure Figure 2J.

### 2.2. Characterization of GelMA-fibrin (GF) hydrogels and assessment of the extent of formation of capillary-like network

GelMA was synthesized using methacrylic anhydride (MA) to enable photo-crosslinking (**Figure 3A**).^[33]^ To verify the percentage of functionalized methacrylation groups, ^1^H NMR was used to measure the extent of free amine group substitution (Figure S2, Supporting Information). The results demonstrated 70% methacrylation of gelatin. Previous studies have shown that GelMAs have integrin-binding motifs and matrix metalproteinase sensitive groups for cells adherence and migration.^[34–35]^ Several successful attempts have been made in generation of functional vascular networks using GelMA.^[36–38]^ On the other hand, fibrin gels have exhibited better angiogenic sprouting and capillary lumen formation with human umbilical vein endothelial cells (HUVECs).^[39–40]^ In order to combine the optimal printing properties of GelMA and the angiogenic properties of fibrin, we used cell-laden inks composed of GelMA and fibrin blends (Figure 3B). Specifically, these materials form a GelMA-fibrin matrix crosslinked by dual-crosslinkers. Thrombin is used to rapidly polymerize fibrinogen, whereas lithium phenyl-2,4,6-trimethylbenzoylphosphinate (LAP) as a photo-initiator for GelMA crosslinking. There are three steps in the formation of the GelMA-fibrin matrix. First, GelMA and fibrinogen undergo a liquid-to-gel transition as the temperature decreases. Second, after UV irradiation exposure, the GelMA is rapidly crosslinked into a gel. The third step involves fibrinogen polymerization into fibrin to form GelMA-fibrin matrix upon submerging into thrombin solution.

**Figure 3.**
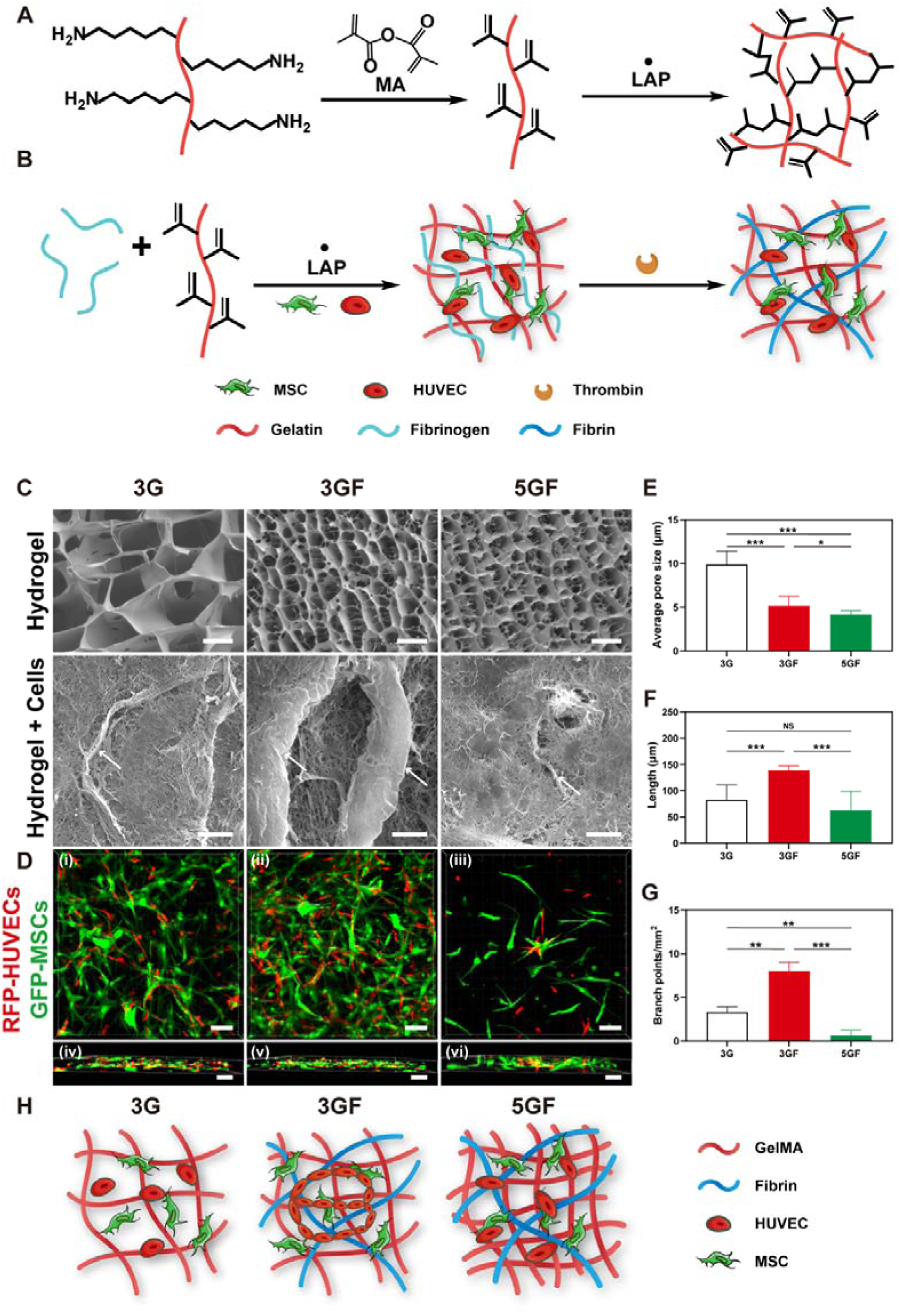
Characterization of GF hydrogels and results depicting the extent of capillary-like network formation. A, B) Schematic diagram depicting GelMA synthesis (A), and steps involved in cell-laden ink, GF ink formation (B). C) SEM images of GelMA or GelMA-fibrin hydrogels with different compositions bearing RFP-HUVCEs and GFP-MSCs. Scale bar represents 10 μm. D) Representative (i-iii) 3D reconstruction of constructs and (iv-vi) Z-plane cross-section from confocal microscopy images showing capillary-like network formation. Scale bar represents 100 μm. E, F) Quantitative analysis of the length (E), and branching points (F) in capillary-like network. G) Graphical representation of morphological parameters of HUVECs and MSCs in different percentages of GelMA-Fibrin IPN. The data are presented as the mean ± SD. *: p<0.1, **: p<0.01, ***: p<0.001.

The microstructures of the hydrogels were observed using scanning electron microscopy (SEM). Micrographs of the GelMA-fibrin blends at different percentage ratios after incubation at 37°C for 3 days have been presented in Figure 3C. As expected, after adding fibrin, the matrix formed a crosslinked interpenetrating polymer network (IPN).^[41]^ To ensure the appropriate biocompatibility, 3% GelMA (3G) and 5% GelMA (5G) were selected as the base, and different concentrations of fibrin solutions were introduced into this base solution. The SEM micrographs revealed that the average size of interconnected pores increased with lowering of the GelMA concentration (Figure 3C). The average size of pores increased with lowering of the concentration of the GelMA used, and decreased after the addition of fibrin (Figure 3E). The porous structure may facilitate nutrient diffusion and improve cell survival. Besides, fibrin fibers were interpenetrating the structure of the GelMA.

Previous studies have shown that the presence of MSCs is crucial for the formation of capillary networks.^[36]^ It was found that the presence of MSCs induced the HUVECs to form HUVEC-networks. To assess the appropriate biocompatibility, RFP-HUVECs and GFP-MSCs were co-encapsulated in a GF matrix and the extent of capillary-like network formation was quantified using confocal microscopy (Figure 3D). Those images clearly reveal that 3% GelMA+0.25% fibrin (3GF) has higher total network length and number of branch points than 5% GelMA+0.25% fibrin (5GF), and 3% GelMA+1% fibrin (3G+1F), respectively (Figure 3F, G; Figure S3 and Movie S1, Supporting Information). These HUVEC-networks were well established, with the majority of them forming cord-like structures. Three-dimensional confocal reconstructed images showed the presence of MSCs adjacent or proximal to the capillary structures, suggesting that the MSCs were differentiating into perivascular cells. It also indicates that 3G is more biocompatible than 5G. Moreover, 3% GelMA consistently generated more robust, interconnected vascular networks with 0.25% fibrin than with 1% fibrin. The formed capillary-like structures in hydrogel were further investigated by SEM. The formed capillary-like structures in 3GF were found to be more abundant and longer than those in 3G or 5GF, which reinforced the findings of the confocal microscopy. Besides, the fiber structure of the hydrogel near the capillary was partially destroyed, which may be due to the degradation of the basement membrane and extracellular matrix by the secreted matrix metalloproteinase (MMP). Hence, 3GF was chosen to be the cell-laden ink of choice pursued in the remaining part of the study. The schematic diagram in Figure 3H presented GelMA and fibrin crosslinked IPNs is based on the obtained results, and depicts HUVECs and MSCs in hydrogels of different compositions.

### 2.3. Tissue perfusion culture and aggregate printing for vascular network formation *in vitro*

In order to provide stable perfusion for long-term culture, we fabricated a continuous flow perfusion system (**Figure 4A**; Figure S4A, Supporting Information). The system includes an incubator, a built-in digital-control peristaltic pump, and a reservoir of culture medium. The designed perfusion chamber (Figure 4B) consisted of three components: a flow part and top and bottom parts. The first one was used for fixing needles, and the top and bottom parts were made of glass which can hold the printed tissue and enabled real-time observation. Before printing, the top and flow parts were assembled respectively, and the flow part contained needle holes for establishing connection with the media perfusion system.

**Figure 4.**
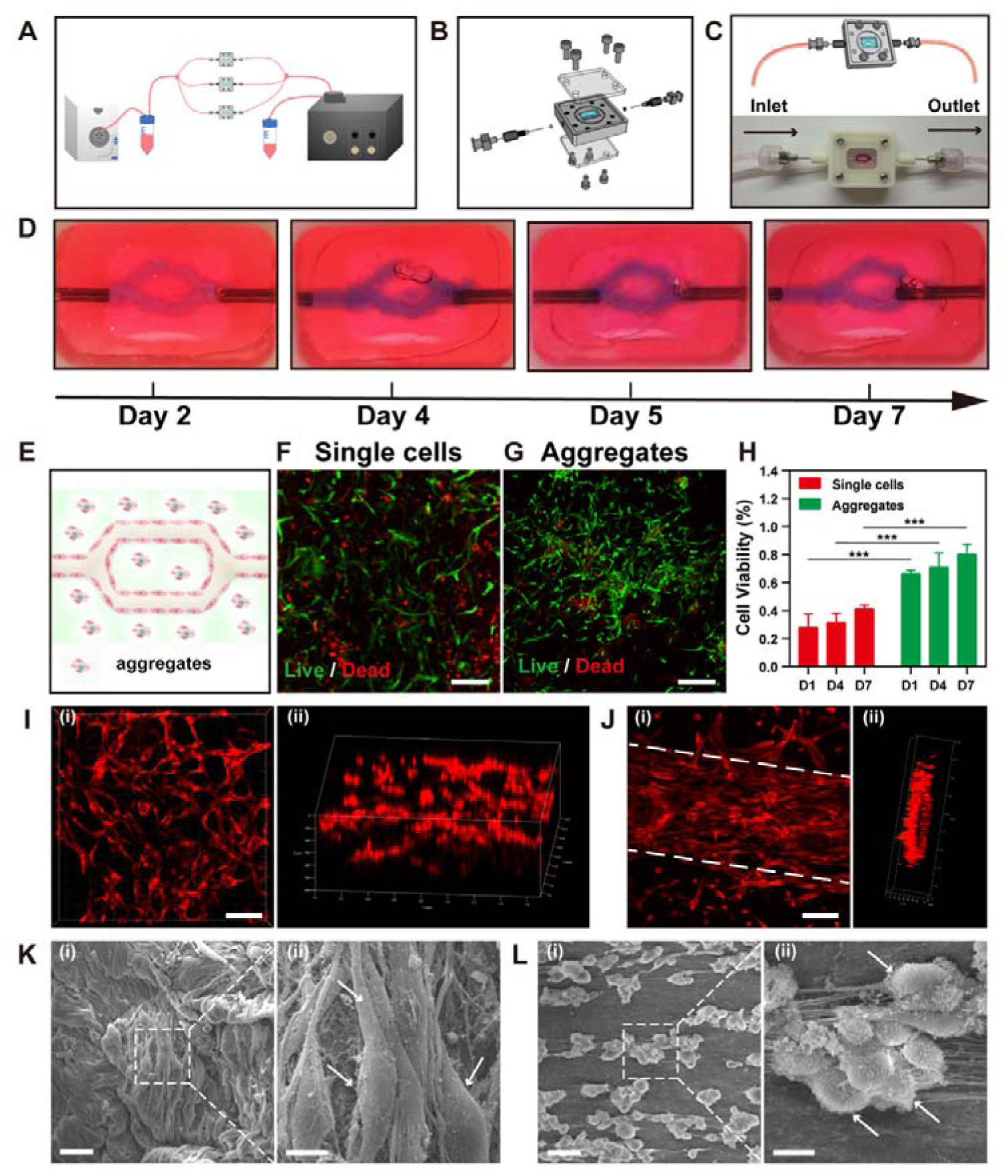
Tissue perfusion culture and aggregate printing for vascular network formation *in vitro*. A) Schematic diagram of perfusion equipment showing digital-control peristaltic pump, reservoir of culture medium, and perfusion chambers containing printed tissues. B) Perfusion chamber schematic diagram showing flow parts, top and bottom glass, luer connector, needles, screw, and sealing ring. C) Schematic diagram showing flow chamber and PU tube-based connections. D) VOs perfused in one week. Images show channel perfusion on days 2, 4, 5, and 7, respectively. E) Schematic diagram showing HUVEC and MSC co-culture aggregates mixed with GF inks printed as VOs. F) Live/dead fluorescence images showing HUVECs and MSCs co-culture aggregates. Scale bar represents 200 μm. G) Live/dead fluorescence images showing co-printed HUVECs and MSCs. Scale bar represents 200 μm. H) Cell viability assay results of printed HUVECs and MSCs co-printed compared with HUVECs and MSCs co-culture aggregates on days 1, 4 and 7 after printing. I) Fluorescence composite images of (i) top view and (ii) cross-section displaying vascularized cells in printed GF inks. Scale bar represents 200 μm. J) Fluorescence composite images of (i) top view and (ii) cross-section displaying RFP-HUVECs seeded in printed channels and perivascular cells. Scale bar represents 200 μm. K) SEM images showing HUVECs and MSCs co-cultures in GF inks after printing. (i) Scale bar represents 20 μm. (ii) Scale bar represents 10 μm. L) SEM images showing HUVECs attached in channels. (i) Scale bar represents 20 μm. (ii) Scale bar represents 10 μm. The data are presented as the mean ± SD. *: p<0.1, **: p<0.01, ***: p<0.001.

To confirm tissue culture capability, we co-printed cell-laden fugitive inks with simple “one-to-two” channeled tissue as described above, with the hydrogels being printed on a specially-designed flow chamber. A 10% GelMA solution was prepared at 37□ and used as elastic ink to fully encapsulate the printed features in the flow chamber, and this was followed by photopolymerization to cross-link the GelMA. Next, the fugitive inks were liquefied and removed from the 3D construct. The flow chamber connected with inlet and outlet tubes through luer connector perfusion system with pumps. Red pigment was added to the medium for coloration (Figure 4C). We used 1.6-mm-inner-diameter tubes and set the inlet perfusion rate at 2-20 μL min for stable and perfusable culture. Importantly, over the course of 7 days of perfusion, the channels always maintained an integrity pattern (Figure 4D; Figure S4B and Movie S2, S3, Supporting Information).

For developing viable strategies for vascularization in tissue engineering, we must carefully explore cell morphology for ensuring high biocompatibility. While cell aggregate cultures hold tremendous potential due to their organotypic cellular interaction, it is easy to improve their viability and function when compared with single cells.^[42–45]^ A previous study has reported the capability of fibroblast aggregates to induce vascularization.^[24]^ We developed aggregates prepared by co-culturing 80% HUVECs and 20% HFFs. It was observed that the HUVECs in the co-culture aggregates assembled to form a vessel-like structure. To fabricate more engineered and vascularized tissues, we used RFP-HUVECs and HFFs co-culture aggregates encapsulated in 3GF inks with gelatin as the fugitive inks. As indicated previously, we initially fabricated the “one-to-two” channeled tissues with HUVECs and MSCs (VOs) shown in Figure 4E. Also, HUVEC and MSC as single cell were encapsulated in 3GF and printed for comparison. To investigate the viability of both single cells and cell aggregates in perfusion culture after 1 days, 4 days and 7 days, we used calcein AM and EthD for staining and confocal imaging, while Image J was used for statistical analysis of data (Figure 4F, G, H; Figure S5, Supporting Information). At day 1, the cell viability was 66% for aggregates. However, the values increased to 71% and 80% at day 4 and day 7, respectively. It was found that printed single cells had no more than 40% viability after 7 days. We found that the initial cell viability was lower compared with that at day 7, suggesting that the printed cells proliferated over time. The observations suggest that our 3D bioprinting approach to printed HUVEC aggregates is less destructive than that observed in single cell.

To demonstrate the formation of vascularized tissues, we used HUVECs and HFFs co-culture aggregates. After a 3-day culture of printed cellular aggregate incubation with 3GF hydrogels, morphological assessment for vascularization and lumen formation was performed (Figure 4I, Movie S4, Supporting Information). Additional network vascularization was studied using 3D-reconstructed confocal, whereby the printed tissues showed multiple cellular aggregates forming capillary-like networks. This observation suggests that those aggregates were important for the formation of vascularized networks *in vitro*. Furthermore, to form vasculature channels, we lined HUVECs in the printed channels of VOs. After overnight perfusion, HUVECs attached to each vessel (Figure 4J). Moreover, it formed a confluent monolayer after one week of culture incubation. It is important that angiogenic HUVEC-based invasions occurred in the vascular channel, with sprout formation bearing lumen-like structures. Although the morphology of the budded sprouts is in its preliminary stages, the exciting perspective is the formation of connections between sprout tips and the HUVEC aggregates. We have demonstrated using SEM the morphology of HUVECs and MSCs encapsulated in 3GF, whereby the attached HUVECs and HUVEC-based aggregates have been shown after different time periods of incubation (Figure 4K, L).

### 2.4 Fabricating vascularized liver tissues as *in vitro* model and its implantation *in vivo*

Previously, it has been shown that a vascularized tissue environment is preferred due to its better cell-cell interactions, and improved hepatocytic function. In this study, in order to fabricate liver models *in vitro*, we encapsulated HAs by replacing HUVEC-based aggregates within the cell-laden hydrogel. HepG2 is a cancer cell line, which has been used as a model cell for tissue regeneration *in vitro* for studying hepatic function.^[46–48]^ For HepG2 aggregate formation, 60% HepG2, 30% HUVECs, and 10% HFFs were mixed, and this was based on the percentages of the different cell types as observed *in vivo*. To confirm the distribution of the three cell types throughout the aggregate, their 3D distribution was assessed using confocal fluorescence microscopy (**Figure 5B**). Notably, after 2 days of incubation, the aggregates’ diameters were about 200 μm and showed albumin (ALB) expression (Figure 5B). Importantly, we printed multi-HAs as shown in Figure 5C, the ALB expression increased after 7 days of perfusion culture *in vitro*. This difference was possibly due to better nutrition and oxygen support in the tissue.

**Figure 5.**
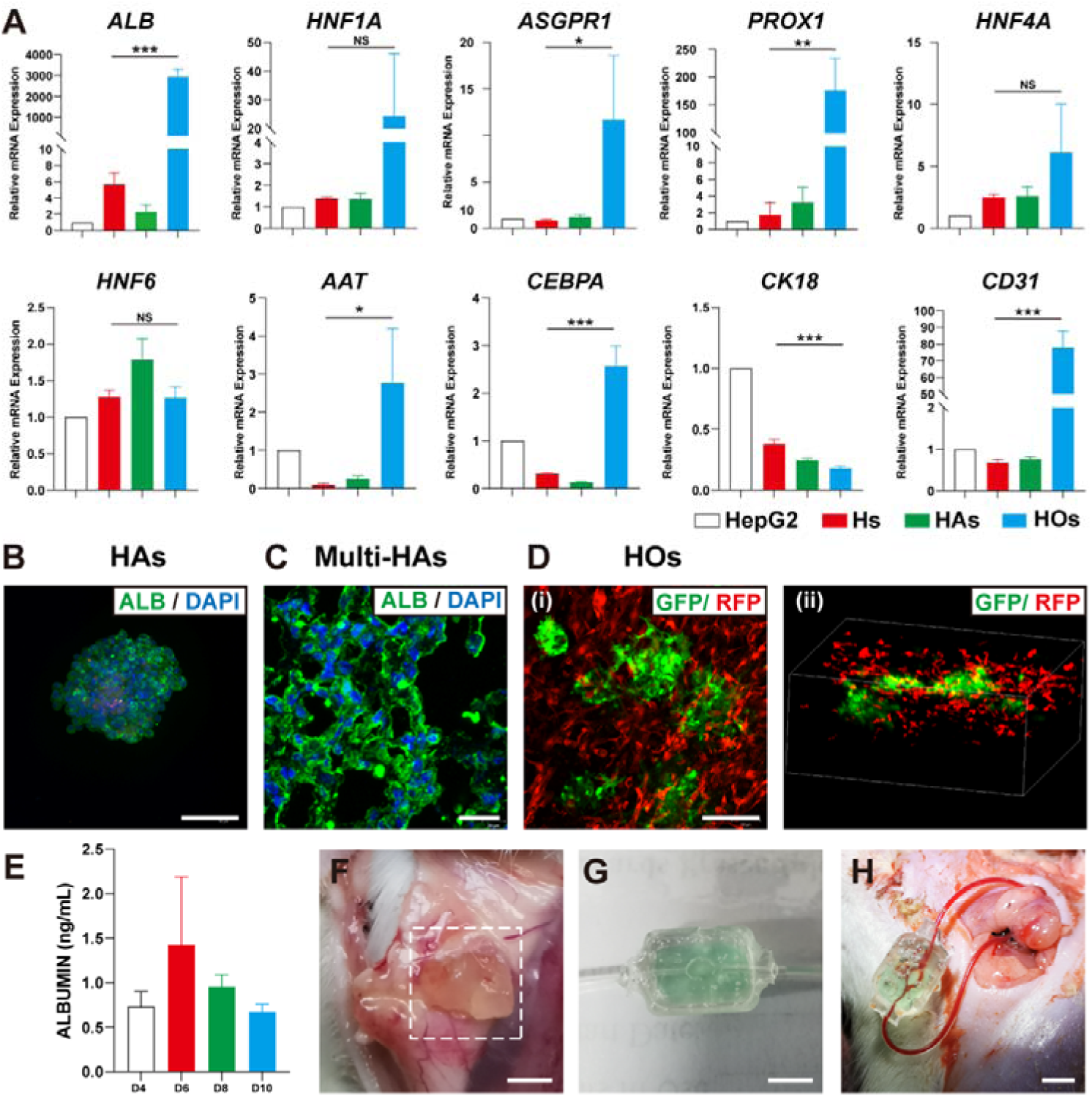
*In vitro* and *in vivo* fabrication of vascular liver. A) Quantitative PCR analysis of the hepatic markers (*ALB*, *HNF1A*, *ASGPR1*, *PROX1*, *HNF4A*, *HNF6*, *AAT*, *CEBPA*, *CK18*), and the vascular marker (*CD31*). B) Fluorescence images of the HAs and multi-HAs showing ALB expression. Scale bar represents 100 μm. C) Fluorescence images of multi-HAs showing ALB expression. Scale bar represents 50 μm. D) Fluorescence images of HOs displaying HAs formed by GFP-HepG2, RFP-HUVECs, and RFP-HUVECs vascularization. Scale bar represents 200 μm. E) ALB expression in cell culture supernatants of HOs as measured by ELISA. F) Observation of transplanted tissues under the subperitoneal zone showing neovascularization of the liver tissue transplants. Scale bar represents 1 cm. G) Images showing “one-to-two” channeled tissues encapsulated in PDMS connected with PU tubes. Scale bar represents 1 cm. H) Observation of transplanted tissues after establishment of connections with carotid artery and jugular vein showing arterial clip slip and establishment of blood perfusion. Scale bar represents 1 cm. The data are presented as the mean ± SD. *: p<0.1, **: p<0.01, ***: p<0.001.

It is observed that HUVECs in multi-HAs do not form connections with each other. Thus, we encapsulated HUVECs, MSCs, and HAs into the matrix hydrogel to generate more vascularization in the tissue. Here, the HUVECs and MSCs were mixed in appropriate ratios. HUVECs and MSCs were seeded at 1×10^7^ cells mL^-1^ and 1×10^6^ cells mL^-1^, respectively. Judging from the 3D-reconstructed confocal images, the printed tissues showed formation of capillary-like networks by HUVECs and MSCs along with formation of multiple cell-aggregates (Figure 5D, Movie S5, Supporting Information). This suggests that HUVECs and MSCs single cells, as well as HAs are important for the formation of vascularized liver tissues *in vitro*. To evaluate the liver function of this tissue, we assessed the expression of hepatic specific genes by quantitative PCR (qPCR) analysis at different periods of culture (Figure S6). It is notable that *ALB* and hepatocyte nuclear factor 4 alpha (*HNF4A*) expression levels were increased and reached peak values after 2 weeks of culturing, and the human alpha 1-antitrypsin (*AAT*) expression was at its highest level at day 4. However, *ASGPR1* and *CK18* expression levels decreased, while the liver tissue *CK18* expression level was lower than that in two-dimensional (2D) culture of HepG2 cells. Compared with 2D HepG2 culture, only HepG2 (Hs) and HAs are printed in the tissue, while the HO expressed the highest levels of *ALB*, *ASGPR1*, *PROX1*, *AAT*, *CEBPA* and *CD31* (Figure 5A). Liver function-related protein expression of ALB between different culture periods was assessed using ELISA (Figure 5E). ALB secretion was the highest after culturing for 6 days, gradually declining thereafter. Thus, liver tissue cultured for 7 days was chosen for further experiments.

To analyze the functionality of vascularized liver tissues *in vivo*, we implanted HOs subperitoneally in mice. We chose 3GF hydrogel as the control. To avoid immunological rejection, after liver tissue transplantation, we injected cyclosporine into the abdominal cavities of the mice. The blood vascular system is a critical system for nutrients and oxygen supply. The liver tissue transplants showed neovascularization at day 7 after transplantation (Figure 5F). However, this phenomenon could not be observed in the 3GF hydrogel treated test group.

To demonstrate that these tissues can be used for direct surgical anastomosis to host vasculature, we leveraged this “one-to-two” channeled tissues combining 5% GelMA as inner elastic inks and external elastic inks outside fugitive inks and cell-laden inks respectively, encapsulated in poly(dimethylsiloxane) (PDMS) were connected to the arteria vessel of adult SD rats, in artery-to-vein mode. The tissues were connected using PU tubes after PDMS-based encapsulation (Figure 5G, Movie S6, Supporting Information). These inlet, outlet PU tubes were connected to the carotid artery and jugular vein, respectively (Figure 5H, Movie S7, Supporting Information). The PU tubes have been shown to be antithrombotic in *in vivo* vascular grafts. To prevent blood clotting in the PU tubes, the animals were injected with heparin during surgery, as well as within 4 days after the surgery. After the vasculature connection being established, the arterial clip was removed to allow blood perfusion. Although this mode was technically challenging owing to the high pressure, the “one-to-two” channel in the hepatic tissue was maintained. One week after the implantation, thrombus was observed and the tissue was removed.

In this study, we present a facile multi-materials bioprinting strategy for constructing centimeter-scale liver-like tissues with branched perfusable vascular networks using a model cell-laden hydrogel formed with organoids bearing bioink. The cell aggregates were deposited in the cell-laden matrix to modulate morphogenesis in space and time. Besides the bioprinted vasculatures, 3D capillary networks were formed successfully using vascularized HUVECs, which supported the nutrient and oxygen requirement of the centimeter-scale liver. We also characterized the HepG2 cells in the tissue, which displayed increased liver-specific gene expression, and determined liver functions, such as albumin secretion, 7 days after differentiation in both *in vitro* and *in vivo* systems. Soft cell-laden elastic hydrogel was tested *in vivo* by direct surgical anastomosis, which has been rarely reported earlier. This work provides a viable and rapid design strategy for biofabrication of engineered tissues, as well as *in vivo* testing using surgical anastomosis for establishing an active vascular network for optimal transport of blood and tissue function.

The printability and biocompatibility of bioink largely affect the construction of soft tissues with vascular network.^[49]^ High concentrations of polymer to some extent can strengthen the hydrogel viscosity.^[50]^ High viscosity allows extruded bioink to better hold its shape and improves mechanical stability, which is especially beneficial in printing larger structures with good resolution.^[51]^ However, higher viscosity increases shear stress during printing, which can damage cells by directly disrupting cell membranes, thereby reducing the rate of proliferation in the surviving cells. Furthermore, it is necessary to use low-concentration (i.e. <5%) GelMA for bioprinting, which can not only achieve higher cell viability after printing, but also facilitate cell proliferation and migration within the printed structure.^[32]^ In the printing process of low-concentration GelMA, problems may occur, including low printing resolution, poor shape fidelity, even nonuniform cell distribution and cell deposition. It’s been a heated research topic on the bioprinting of low-concentration (i.e. <5%) GelMA. Current approaches adapted to tackle the challenge mainly include the additives of other materials and printing hydrogels under low temperature.^[32, 52]^ Technically speaking, adding other materials is not categorized in the field of low-concentration bioprinting. What’s more, bioprinting hydrogel under low temperature may be problematic including poor shape fidelity and decreased cell viability (nearly 90%). Therefore, low-concentration GelMA could achieve fine bioprinting with enhancing high cell viability, which is a significant research problem to be resolved. In this study, we used 3% GelMA as cell-laden inks with the secondary temperature of the BiopHead, which promote high cell viability and printing accuracy.

The vasculature is responsible for mass transport which sustains the metabolic requirements of engineered tissues. Dynamic culture under continuous perfusion could supply nutrients and oxygen, while removing metabolic wastes through stable inter-connected vascular networks.^[53–54]^ In continuous perfusion culture such as cell aggregates and organoids, higher cell densities can be sustained.^[20]^ Furthermore, this versatile platform also can be used to precisely control the growth and differentiation of cultured tissue.^[17]^ With addition of appropriate growth factors and culture medium, perfusion tissue can be stimulated to attain maturity and achieve efficient functionalization.^[55]^

## 3. Conclusions

Three-dimensional printing holds great promise for engineering whole organs, but the field is still in its nascent stage with many unresolved challenges. In this study, we demonstrate the use of low concentrations of GelMA with fibrin for the first time, as a bioink for extrusion-based printing of vascularized tissues, producing model organ tissues with high-resolution and high cell viability. This 3D engineering-based methodology has allowed us to fabricate vascular networks in centimeter-scale tissues with active tissue functionality. Furthermore, it opens a new avenue for fundamental drug screening studies.

## 4. Experimental Section

### GelMA synthesis

We used a simple method to synthesize GelMA. First, gelatin (Type A, 300 bloom from porcine skin, Sigma Aldrich) was dissolved in Dulbecco’s phosphate-buffered saline (DPBS, Gibco) to make a 10% (w/v) solution, and stirred at 60°C for 2 h. Then the temperature was lowered to 50°C, and 0.5 ml methacrylic anhydride (Sigma Aldrich) was slowly added to the gelatin solution with continuous stirring at a rate of 1 mL min^-1^ per gram of gelatin, to obtain 70% modified GelMA. After 3 hours of reaction, the temperature was reduced to 40°C, and the reaction solution was dialyzed with deionized water. The molecular weight cut-off of the dialysis bag was 14 kDa (Spectrum Labs, Inc.). In order to remove excess methacrylic acid and salts, dialysis was performed at 40°C, and the fluid was changed once a day for a week. The reaction solution was aliquoted, lyophilized, and stored at −80°C. The lyophilized powder was stored at -20°C. Pre-heated sterile DPBS at 60°C was used to dissolve 10% GelMA, and 1 M NaOH was used to adjust the pH to 7.2-7.4. The methacrylation degree of free amine group in GelMA sample was determined by ^1^H NMR as previously described.^[56]^ Before use, 0.1% (w/v) lithium phenyl-2, 4, 6-trimethylbenzoylphosphinate (LAP, Sigma Aldrich) was added.

### Bioink preparation

We prepared several types of inks for 3D printing. Cell-laden inks were prepared using GelMA and fibrin (Sigma Aldrich, MO, USA) at different mixing ratios. Fibrinogen was stored at concentration of 25 mg mL^-1^ in sterile DPBS without calcium and magnesium, and kept at 37°C for 30 min to allow full dissolution. Photocrosslinking was achieved by exposing the GelMA or GelMA-Fibrin prepolymer to 10 mW cm^-2^ UV light (365 nm, Goodun) for 2 min.

The 10% gelatin-based bioink was used as the sacrificial material, which was dissolved in pre-heated sterile DPBS at 60°C, and the pH was adjusted to 7.2-7.4 with 1 M NaOH. A 10% GelMA solution was used as the elastic material for the *in vitro* perfusion culture. The elastic material used for jugular artery and vein transplantation was poly(dimethylsiloxane) (PDMS, Sylgard^TM^ 184 silicone elastomer kit, Dow chemical company). This ink is composed of two elastomers in a weight ratio of 10:1, which were mixed for 30 s at 2000 rpm with a stirring mixer (AE-310, Thinky Corporation) to make the final solution uniform.

### Viscosity test

A viscometer (RST-CPS cone plate rheometer, Brookfield) was used to measure the viscosity of 1%, 2%, 3%, 4%, 5% GelMA, and 3% GelMA+0.25% Fibrin bioink, respectively. First, the temperature of the viscometer measuring plate with a upper plate diameter of 25 mm was cooled to 2□. The GelMA solution was dropped on the measuring plate, and the distance between the plates was adjusted to 50 μm. The excess material was wiped off and allowed to stand for 1 min. The upper measuring plate (rotor) was gently rotated before measurement to avoid adhesion between the plates, and the viscosity was measured at a shear rate of 10 s^-1^ with temperature increase of 1°C per min.

### Morphology and porosity analysis by Scanning Electron Microscopy (SEM)

The GelMA hydrogel and GelMA-Fibrin hydrogel were immersed in PBS for 24 h at 37□. For studies of internal porosity, samples were fixed with 3.7% paraformaldehyde (PFA, Fluka) for 30 min, immersed in liquid nitrogen for 60 seconds, freeze-fractured using a cold razor blade, and sublimation at -75□ for 45min, and then sputter coated with Au. All samples were observed by HITACHI S-3000N&Quorum PP3000T Scanning Electron Microscope.

### SEM images for hydrogel with cells

the hydrogel samples with cells were soaked with 2.5% glutaraldehyde at 4□ for 24 h. Then all samples were washed with PBS for 3 times, and dehydrated samples through graded ethanol solutions with 15, 30, 50, 70, 80, 90, 100, 100% (v/v, in water) for 10 min each time. After that all samples were dried by a critical point drying (CPD) technique using a Bal-Tec 030 instrument. Finally, all samples were coated with Au by a sputter and examined with SEM (phenom pro).

### Cell culture and maintenance

Primary human umbilical vein endothelial cells (HUVECs) and red fluorescent protein-expressing HUVECs (RFP-HUVECs) were maintained in EMG-2 medium (complete EGM-2 BulletKit^TM^, Lonza). We obtained human umbilical cords from Xinhua Hospital Affiliated to Shanghai Jiaotong University School of Medicine. The collection and use of the obtained umbilical cords were approved by the institutional ethical committee (approval number: XHEC-C-2020-092-1), and informed consent was obtained from all participants. HepG2, human foreskin fibroblast (HFFs), and human umbilical cord MSCs were donated by the National Stem Cell Resource Center, Beijimg. HepG2, GFP-HepG2 and HFFs were cultured in Dulbelco’s modified Eagle medium containing high glucose and sodium pyruvate (DMEM, Gibco) supplemented with 10% fetal bovine serum (FBS, Bioind), 1% penicillin/streptomycin (Gibco). MSCs and GFP-MSCs were cultured in Dulbecco’s modified Eagle medium containing high glucose and sodium pyruvate (DMEM, Gibco) supplemented with 15% fetal bovine serum (FBS, Bioind), 1% non-essential amino acid solution (NEAA, Gibco), 1% GlutaMAXTM (Gibco), 1% penicillin/streptomycin (Gibco). All cells were cultured at 37°C and 5% CO_2_ in an incubator, and the medium was changed every 2 days. HUVECs, RFP-HUVECs and HFFs were not used beyond the 10^th^ passage.

For HUVEC cell aggregate preparation, 8×10^4^ HUVECs and 2×10^4^ HFFs were suspended in 1.5 mL medium and seeded into each well of a 24-well Kuraray ultra-low attachment plate (round-bottom type, Elplasia). Each plate has 400 microwells. Cells supplemented with EGM-2 were seeded evenly in the microwells, which were then allowed to self-aggregate over a time of 24□h. For HepG2 aggregate preparation, 6×10^4^ HepG2, 3×10^4^ HUVECs, and 1×10^4^ HFFs were suspended in 1.5 mL medium and seeded in a well. Cells were maintained in EGM-2, and the medium was changed 2-3 times every day by replacing 1 mL of the supernatant in each well.

### Encapsulation of HUVECs and MSCs in hydrogels

When cells cultured reached 90% confluency, the culture medium was discarded and cells were washed with PBS, and then incubated with 0.25% Trypsin-EDTA (Gibco) for 1 min at 37°C to detach the cells from the culture dishes. The cell suspension was centrifuged at 1200 rpm for 3 min at room temperature. The supernatant was discarded, and the cells were resuspended in hydrogels at 37 °C. For RFP-HUVECs and GFP-MSCs co-encapsulation, we mixed 1×10^6^ RFP-HUVECs and 1×10^6^ GFP-MSCs with 3% GelMA, 3% GelMA+0.25% fibrin, 3% GelMA+1% fibrin, 5% GelMA, 5% GelMA+0.25% fibrin, respectively. A volume of 20 μL of the cell-prepolymer mixture was dispensed in each well of a 24-well flat bottom cell culture plate. Photocrosslinking was achieved by exposing the mixture to 10 mW cm^-2^ UV light (365nm, Goodun) for 2 min. The encapsulated hydrogels were then cultured with EGM-2 medium.

### Bioprinting platform and hardware

A 3D bioprinter developed by the Shenyang Institute of Automation, Chinese Academy of Sciences, was used for the 3D vascular tissue printing experiment. The main components of the printer include 5 print heads, a print platform, and a print bin, which will allow multi-materials printing. As the core component of the printer, the print head included an integrated electrical interface, a two-stage temperature control unit, a two-stage temperature sensor, a water-cooling block, and two printing components (electric extrusion and pneumatic extrusion). The printing platform can be set at a wide range of temperatures ranging from -10°C to 60°C. The printed 3D vascular network tissue was cured using a low-temperature gel. The printing chamber has an ultraviolet sterilization function which allows aseptic printing.

### Fabrication process

The 3D drawing software SolidWorks 2018 (SolidWorks Software, Inc., La Jolla, USA) was used to model the designed vascular tissue, import the geometric model into the self-developed software Bipcoder to configure printing parameters, and generate the G code. For *in vitro* perfusion culturing, 10% gelatin, 3% GelMA+0.25% fibrin were used as the sacrificial material and the cell-laden material respectively. The feeder needles transfer those inks into the two print heads, and allow maintenance of the temperature of the printing silo and the printing needle, respectively. The temperature of the printing silo was set slightly higher (8°C) than the gel point temperature (6°C), and the temperature of the printing needle was based on the temperature representing the gel point in the printability map. For transplantation printing, 10% gelatin, 3% GelMA+0.25% fibrin, and 5% GelMA were used as the sacrificial material, cell-laden material, and elastic material, respectively, and loaded into respective print heads. All needles used for printing have a diameter of 200 μm. Photocrosslinking was achieved using UV light (365nm, Goodun) at the wavelength of 365 nm for 2 min after printing was completed. Then it is placed in thrombin solution (5 U mL^-1^) at 37°C to dissolve and remove the internal gelatin material and allow formation of a vascular channel structure.

### Endothelial monolayer and vascular network formation

The printed structure was mounted in a customized chamber. We used 10% GelMA to encapsulate the printed constructs for perfusion. The cells were resuspended to have 1 × 10^7^ cells mL^-1^ in EGM-2 medium. To form an endothelial monolayer, 10 μL cell resuspension was seeded into the channel using a micropipette. Adhesion of the cells to the bottom surface was allowed for 30 min, followed by flipping of the system and incubation for 30 min to allow adherence to the top surface. After the seeding period, the constructs were put into the perfusion incubator (TEB500, Ebers) with the tubing inset into the flow chamber. Unattached cells were washed away with culture medium EGM-2 and 5 ng mL^-1^ VEGF (R&D). Perfusion rate was set to 2-20 μL min^-1^. The constructs were cultured for several days before further characterizations.

### Cell viability assay

Cell viability was analyzed using LIVE/DEAD Viability/Cytotoxicity Kit (Life Technologies). After washing the cells three times with phosphate-buffered saline (PBS), a PBS solution containing 0.5 μL mL calcein AM and 2 μL mL EthD was used to stain the cells for 20 min in the dark. The cells were then washed three times with PBS to remove residual regents. An inverted confocal microscope was used to take the fluorescence images (Calcein AM: Ex 488 nm, Em 505-525 nm; EthD: Ex 559 nm, Em 600-630 nm).

### SD mice and HOs transplantation

All animal experiments were approved by the Institutional Animal Care and Use Committee (IACUC) of the Institute of Zoology, Chinese Academy of Sciences in accordance with institutional and national guidelines (Ethical approval No. IOZ20180063). To analyze the functionality of vascularized liver tissues in vivo, we implanted HOs it into subperitoneal of mice after anaesthetising.

### Immunofluorescence staining

For ALB staining, printed tissues were fixed with 4% paraformaldehyde for 30 min at room temperature, and permeabilized with 0.5% Triton X-100 (Sigma) for 30 min, followed by PBS washing (3 times). The tissues were blocked with 3% bovine serum albumin (BSA) in PBS overnight. Then they were incubated for 2 days with a primary antibody anti-ALB (R&D, MAB1455-SP, 1:100) in blocking buffer at 4 °C, followed by PBS washing (3 times). The tissues were then incubated with fluorescence-conjugated secondary antibodies Alexa Fluor 488 (Donkey anti-mouse, Invitrogen, A21202, 1:500) diluted in PBS for 2 days in the dark at 4°C. Finally, the nuclei were stained with Hoechst 33342 (Invitrogen, 1:1000) for 10 min, followed by three PBS washes.

### Real-time quantitative polymerase chain reaction

Total RNA was extracted from printed tissues and cells using TRIzol (Invitrogen), following the manufacturer’s instructions. One microgram of RNA was reverse transcribed into cDNA using a PrimeScript™ RT reagent Kit (TaKaRa, RR037A) in a 20 μl reaction. Real-time quantitative PCR (qPCR) was performed and analyzed using a CFX96^TM^ real-time system (Bio-Rad) with TB Green Premix Ex Taq™ (TaKaRa, RR420A). The expression level of *GAPDH* was used for internal normalization. The details of the primers for HepG2 and HUVECs have been listed in Supplementary Table 1.

### Albumin ELISAs

To measure ALB secretion, the 3D printed tissues were cultured for various time periods. Culture supernatants were collected at 24 h after a medium change and stored at -80°C before analysis. ELISAs were performed using a human albumin ELISA kit (Abcam, ab179887) according to the manufacturer’s instructions. *Imaging and analysis*: Photographs of fabricated tissues were acquired using Leica SAPO stereo microscopes and high-speed CMOS cameras (PCO. dimax HS, PCO). Confocal microscopy was performed using a Leica Dmil fluorescent microscope and a Zeiss LSM 780 fluorescent microscope. Three-dimentional projections were generated in Imaris (Imaris 9.0.2, Bitplane Scientific Software) and ZEN software (Zeiss). For cell counting and length calculation, a semi-automated process in Image J was used.

### Statistics

Data are presented as mean ± SD (n=3). The t-test was performed to determine significant differences using GraphPad Prism 8 (GraphPad Software, Inc., La Jolla, USA). The significance levels (p-values) are indicated with asterisks and specific p-values are provided in each figure legend.

## Supporting Information

Supporting Information is available from the Wiley Online Library or from the author.

## Acknowledgements

Xin Liu, Xinhuan Wang, Lulu Sun, and Liming Zhang contributed equally to this work. The authors acknowledge funding support from the Strategic Priority Research Program of Chinese Academy of Sciences, Grant No. XDA16020802, XDA16020803 and XDA16020804, National Key Research and Development Program of China 2018YFE0204403, CAS Pioneer Hundred Talents Program, Y829F11102, the National Key Research and Development Project under Grant 2020YFB1313100, the National Natural Science Foundation of China (51875557), the Research Equipment Development Program of the Chinese Academy of Sciences (YJKYYQ20170042, YJKYYQ20190045), Foundation of State Key Laboratory of Robotics (2017-Z16), the China Postdoctoral Science Foundation (2020M670454), and Financial supports from the State Key Laboratory of Membrane and the State Key Laboratory of Robotics are also gratefully acknowledged. We thank Mr. Kai Li for technical support of graphical design and Dr. Ji for grammar revision.

## Author Contributions

X. Liu developed the multi-scale vascular tissues, designed and performed experiments, analyzed data and prepared manuscript. Dr. X. Wang contributed to GelMA synthesis, polymer characterization, SEM analysis, endothelization and vascular network formation, subperitoneally transplantation and carotid artery and jugular vein connection. L. Zhang performed viscosity test, bioprinting parameter testing, fabrication process. Dr. L. Sun performed HUVECs and MSCs encapsulation experiments, HUVEC cell aggregate preparation. Dr. H. Wang performed bioprinting platform and hardware designed. Dr. H. Zhao performed GelMA synthesis. Z. Zhang performed the HUVECs isolation. Dr. Y. Huang performed polymer characterization. J. Zhang and B. Song performed perfusion experiments. C. Li performed HUVECs and MSCs encapsulation experiments. H. Zhang and S. Li performed bioprinting parameter testing, fabrication process.

## Conflict of Interest

The authors declare no conflict of interest.

## Date Availability Statement

The data that support the findings of this study are available from the corresponding author upon reasonable request.

## The Table of Contents

The 3D printing engineering method reported in this article fabricated centimeter-scale vascularized soft tissues with high viability and accuracy using multi-materials bioprinting. It allowed us to fabricate vascular networks in centimeter-scale hepatic tissues with active tissue functionality *in vitro* and *in vivo*, which will potentially provide a model for fundamental drug screening studies.

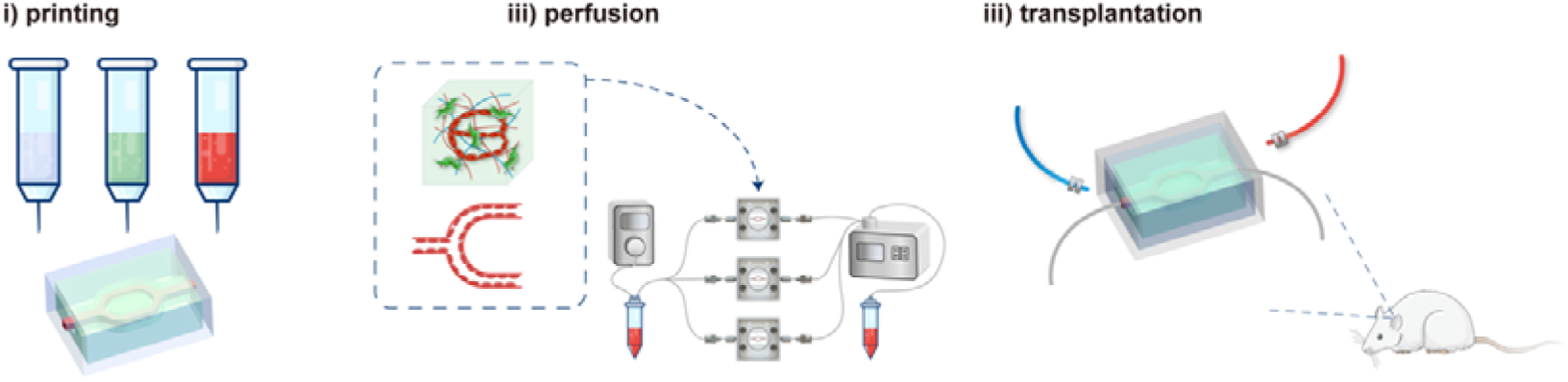

## Notes

### Competing Interest Statement

The authors have declared no competing interest.

## References

1. S. V. Murphy, A. Atala, Nat. Biotechnol. 2014, 32, 773.

2. Y. S. Zhang, K. Yue, J. Aleman, K. M. Moghaddam, S. M. Bakht, J. Yang, W. Jia, V. Dell’Erba, P. Assawes, S. R. Shin, M. R. Dokmeci, R. Oklu, A. Khademhosseini, Ann. Biomed. Eng. 2017, 45, 148.

3. L. Moroni, T. Boland, J. A. Burdick, C. De Maria, B. Derby, G. Forgacs, J. Groll, Q. Li, J. Malda, V. A. Mironov, C. Mota, M. Nakamura, W. Shu, S. Takeuchi, T. B. F. Woodfield, T. Xu, J. J. Yoo, G. Vozzi, Trends Biotechnol. 2018, 36, 384.

4. T. Gao, G. J. Gillispie, J. S. Copus, A. K. Pr, Y. J. Seol, A. Atala, J. J. Yoo, S. J. Lee, Biofabrication 2018, 10, 034106.

5. P. S. Gungor-Ozkerim, I. Inci, Y. S. Zhang, A. Khademhosseini, M. R. Dokmeci, Biomater. Sci. 2018, 6, 915.

6. Y. S. Zhang, A. Khademhosseini, Science 2017, 356, eaaf3627.

7. N. F. Huang, V. Serpooshan, V. B. Morris, N. Sayed, G. Pardon, O. J. Abilez, K. H. Nakayama, B. L. Pruitt, S. M. Wu, Y. S. Yoon, J. Zhang, J. C. Wu, Commun. Biol. 2018, 1, 199.

8. S. Maharjan, J. Alva, C. Cámara, A. G. Rubio, D. Hernández, C. Delavaux, E. Correa, M. D. Romo, D. Bonilla, M. L. Santiago, W. Li, F. Cheng, G. Ying, Y. S. Zhang, Matter 2021, 4, 217.

9. D. Richards, J. Jia, M. Yost, R. Markwald, Y. Mei, Ann. Biomed. Eng. 2017, 45, 132.

10. C. K. Griffith, C. Miller, R. C. A. Sainson, J. W. Calvert, N. L. Jeon, C. C. W. Hughes, S. C. George, Tissue Engineering 2005, 11, 257.

11. Y. S. Zhang, A. Khademhosseini, in Tissue-Engineered Vascular Grafts, DOI: 10.1007/978-3-030-05336-9_11 2020, Ch. Chapter 11, p. 321.

12. X. Cao, S. Maharjan, R. Ashfaq, J. Shin, Y. Shrike Zhang, Engineering 2020, DOI: https://doi.org/10.1016/j.eng.2020.03.019.

13. P. Datta, B. Ayan, I. T. Ozbolat, Acta Biomater. 2017, 51, 1.

14. I. T. Ozbolat, M. Hospodiuk, Biomaterials 2016, 76, 321.

15. H. Gudapati, M. Dey, I. Ozbolat, Biomaterials 2016, 102, 20.

16. J. S. Miller, K. R. Stevens, M. T. Yang, B. M. Baker, D.-H. T. Nguyen, D. M. Cohen, E. Toro, A. A. Chen, P. A. Galie, X. Yu, R. Chaturvedi, S. N. Bhatia, C. S. Chen, Nat. Mater. 2012, 11, 768.

17. D. B. Kolesky, K. A. Homan, M. A. Skylar-Scott, J. A. Lewis, Proc. Natl. Acad. Sci. USA 2016, 113, 3179.

18. L. E. Bertassoni, M. Cecconi, V. Manoharan, M. Nikkhah, J. Hjortnaes, A. L. Cristino, G. Barabaschi, D. Demarchi, M. R. Dokmeci, Y. Yang, A. Khademhosseini, Lab on a Chip 2014, 14, 2202.

19. L. Zhao, V. K. Lee, S.-S. Yoo, G. Dai, X. Intes, Biomaterials 2012, 33, 5325.

20. M. A. Skylar-Scott, S. G. Uzel, L. L. Nam, J. H. Ahrens, R. L. Truby, S. Damaraju, J. A. Lewis, Sci. adv. 2019, 5, eaaw2459.

21. B. Trappmann, B. M. Baker, W. J. Polacheck, C. K. Choi, J. A. Burdick, C. S. Chen, Nat. Commun. 2017, 8, 371.

22. T. J. Hinton, Q. Jallerat, R. N. Palchesko, J. H. Park, M. S. Grodzicki, H.-J. Shue, M. H. Ramadan, A. R. Hudson, A. W. Feinberg, Sci. Adv. 2015, 1, e1500758.

23. H.-H. G. Song, K. M. Park, S. Gerecht, Adv. Drug Deliv. Rev. 2014, 79–80, 19.

24. Y. Nashimoto, T. Hayashi, I. Kunita, A. Nakamasu, Y.-s. Torisawa, M. Nakayama, H. Takigawa-Imamura, H. Kotera, K. Nishiyama, T. Miura, R. Yokokawa, Integrative Biology 2017, 9, 506.

25. I. T. Ozbolat, Trends Biotechnol. 2015, 33, 395.

26. J. D. Baranski, R. R. Chaturvedi, K. R. Stevens, J. Eyckmans, B. Carvalho, R. D. Solorzano, M. T. Yang, J. S. Miller, S. N. Bhatia, C. S. Chen, Proc. Natl. Acad. Sci. USA 2013, 110, 7586.

27. X. Ye, L. Lu, M. E. Kolewe, H. Park, B. L. Larson, E. S. Kim, L. E. Freed, Biomaterials 2013, 34, 10007.

28. B. Zhang, M. Montgomery, M. D. Chamberlain, S. Ogawa, A. Korolj, A. Pahnke, L. A. Wells, S. Masse, J. Kim, L. Reis, A. Momen, S. S. Nunes, A. R. Wheeler, K. Nanthakumar, G. Keller, M. V. Sefton, M. Radisic, Nat. Mater. 2016, 15, 669.

29. I. S. Kinstlinger, S. H. Saxton, G. A. Calderon, K. V. Ruiz, D. R. Yalacki, P. R. Deme, J. E. Rosenkrantz, J. D. Louis-Rosenberg, F. Johansson, K. D. Janson, D. W. Sazer, S. S. Panchavati, K.-D. Bissig, K. R. Stevens, J. S. Miller, Nat. Biomed. Eng. 2020, 4, 916.

30. C. Colosi, S. R. Shin, V. Manoharan, S. Massa, M. Costantini, A. Barbetta, M. R. Dokmeci, M. Dentini, A. Khademhosseini, Adv. Mater. 2016, 28, 677.

31. J. Yin, M. Yan, Y. Wang, J. Fu, H. Suo, ACS Appl. Mater. Interfaces 2018, 10, 6849.

32. W. Liu, M. A. Heinrich, Y. Zhou, A. Akpek, N. Hu, X. Liu, X. Guan, Z. Zhong, X. Jin, A. Khademhosseini, Y. S. Zhang, Adv. Healthc. Mater. 2017, 6.

33. A. I. Van Den Bulcke, B. Bogdanov, N. De Rooze, E. H. Schacht, M. Cornelissen, H. Berghmans, Biomacromolecules 2000, 1, 31.

34. J. W. Nichol, S. T. Koshy, H. Bae, C. M. Hwang, S. Yamanlar, A. Khademhosseini, Biomaterials 2010, 31, 5536.

35. M. Nikkhah, N. Eshak, P. Zorlutuna, N. Annabi, M. Castello, K. Kim, A. Dolatshahi-Pirouz, F. Edalat, H. Bae, Y. Yang, A. Khademhosseini, Biomaterials 2012, 33, 9009.

36. Y.-C. Chen, R.-Z. Lin, H. Qi, Y. Yang, H. Bae, J. M. Melero-Martin, A. Khademhosseini, Adv. Funct. Mater. 2012, 22, 2027.

37. D. B. Kolesky, R. L. Truby, A. S. Gladman, T. A. Busbee, K. A. Homan, J. A. Lewis, Adv. Mater. 2014, 26, 3124.

38. W. Jia, P. S. Gungor-Ozkerim, Y. S. Zhang, K. Yue, K. Zhu, W. Liu, Q. Pi, B. Byambaa, M. R. Dokmeci, S. R. Shin, A. Khademhosseini, Biomaterials 2016, 106, 58.

39. R. R. Rao, A. W. Peterson, J. Ceccarelli, A. J. Putnam, J. P. Stegemann, Angiogenesis 2012, 15, 253.

40. G. A. Calderon, P. Thai, C. W. Hsu, B. Grigoryan, S. M. Gibson, M. E. Dickinson, J. S. Miller, Biomater. Sci. 2017, 5, 1652.

41. A. P. Dhand, J. H. Galarraga, J. A. Burdick, Trends Biotechnol. 2020, DOI: https://doi.org/10.1016/j.tibtech.2020.08.007.

42. A. C. Daly, M. D. Davidson, J. A. Burdick, Nat. Commun. 2021, 12, 753.

43. D. J. Richards, Y. Li, C. M. Kerr, J. Yao, G. C. Beeson, R. C. Coyle, X. Chen, J. Jia, B. Damon, R. Wilson, E. Starr Hazard, G. Hardiman, D. R. Menick, C. C. Beeson, H. Yao, T. Ye, Y. Mei, Nat. Biomed. Eng. 2020, 4, 446.

44. M.-O. Lee, K. B. Jung, S.-J. Jo, S.-A. Hyun, K.-S. Moon, J.-W. Seo, S.-H. Kim, M.-Y. Son, Journal of Biological Engineering 2019, 13, 15.

45. Q. Tan, K. M. Choi, D. Sicard, D. J. Tschumperlin, Biomaterials 2017, 113, 118.

46. H. Jeon, K. Kang, S. A. Park, W. D. Kim, S. S. Paik, S.-H. Lee, J. Jeong, D. Choi, Gut Liver 2017, 11, 121.

47. T. Billiet, E. Gevaert, T. De Schryver, M. Cornelissen, P. Dubruel, Biomaterials 2014, 35, 49.

48. C.-A. E. Suurmond, S. Lasli, F. W. van den Dolder, A. Ung, H.-J. Kim, P. Bandaru, K. Lee, H.-J. Cho, S. Ahadian, N. Ashammakhi, M. R. Dokmeci, J. Lee, A. Khademhosseini, Adv. Healthc. Mater. 2019, 8, 1901379.

49. A. Panwar, L. P. Tan, Molecules 2016, 21.

50. V. H. Mouser, F. P. Melchels, J. Visser, W. J. Dhert, D. Gawlitta, J. Malda, Biofabrication 2016, 8, 035003.

51. N. Paxton, W. Smolan, T. Bock, F. Melchels, J. Groll, T. Jungst, Biofabrication 2017, 9, 044107.

52. J. Yin, M. Yan, Y. Wang, J. Fu, H. Suo, ACS Appl. Mater. Interfaces 2018, 10, 6849.

53. M. Radisic, J. Malda, E. Epping, W. Geng, R. Langer, G. Vunjak-Novakovic, Biotechnol. Bioeng. 2006, 93, 332.

54. A. Tocchio, M. Tamplenizza, F. Martello, I. Gerges, E. Rossi, S. Argentiere, S. Rodighiero, W. Zhao, P. Milani, C. Lenardi, Biomaterials 2015, 45, 124.

55. S. Alimperti, T. Mirabella, V. Bajaj, W. Polacheck, D. M. Pirone, J. Duffield, J. Eyckmans, R. K. Assoian, C. S. Chen, Proc. Natl. Acad. Sci. USA 2017, 114, 8758.

56. D. Wei, W. Xiao, J. Sun, M. Zhong, L. Guo, H. Fan, X. Zhang, J Mater. Chem. B 2015, 3, 2753.

