## Supplementary material for "A novel method for generating 3D constructs with branched vascular networks using multi-materials bioprinting and direct surgical anastomosis": Methods and supplemental figs

Dr. X. Wang, J. Zhang, Prof. Q. Gu

State Key Laboratory of Membrane Biology, Institute of Zoology, Chinese Academy of Sciences, Beijing 100101, P. R. China

X. Liu, Prof. Q. Gu

Savaid Medical School, University of Chinese Academy of Sciences, Beijing 100049, P. R. China

L. Zhang, Dr. H. Wang, Hui Zhang, Song Li, Prof. X. Zheng

Shenyang Institute of Automation, Chinese Academy of Sciences, Shenyang 110169, P. R. China

Dr. H. Zhao, Dr. Y. Huang, Prof. S. Wang

Institute of Chemistry, Chinese Academy of Sciences, Beijing, 100190 P. R. China

Dr. L. Sun, Z. Zhang, C. Li

State Key Laboratory of Cell Biology, CAS Center for Excellence in Molecular Cell Science, Shanghai Institute of Biochemistry and Cell Biology, Chinese Academy of Sciences, Shanghai 200031, P. R. China

B. Song

University of Science and Technology of China, Hefei, 230026, P. R. China

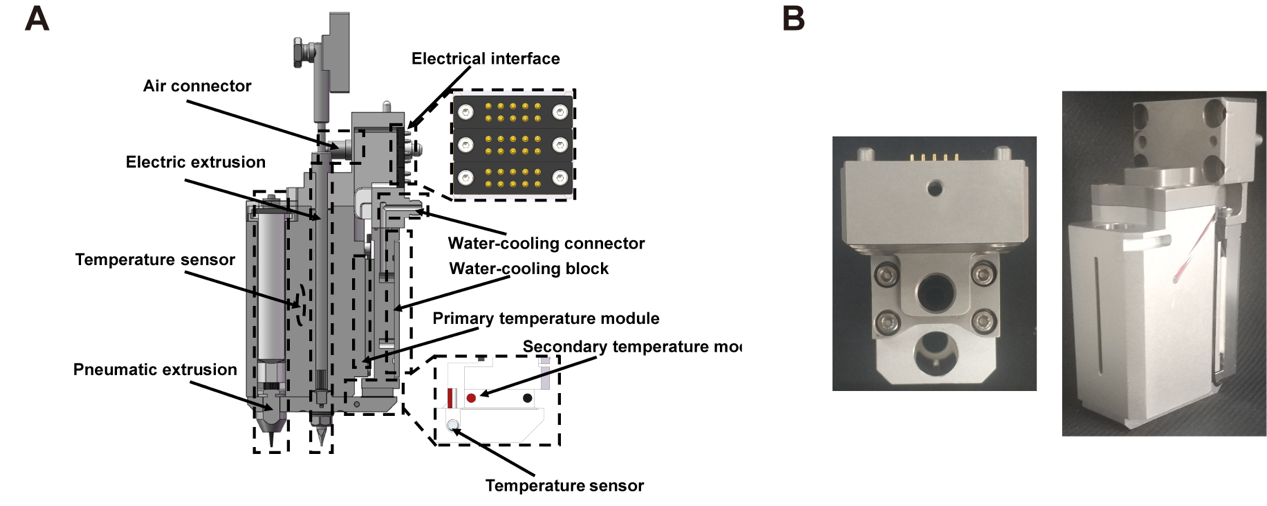

**Figure S1.** A) Graph of BiopHead showing air connector, electric extrusion, temperature senor, pneumatic extrusion, electrical interface, water-cooling connector, water-cooling block, primary temperature module and second temperature module. B) images of side view and top view of BiopHead.

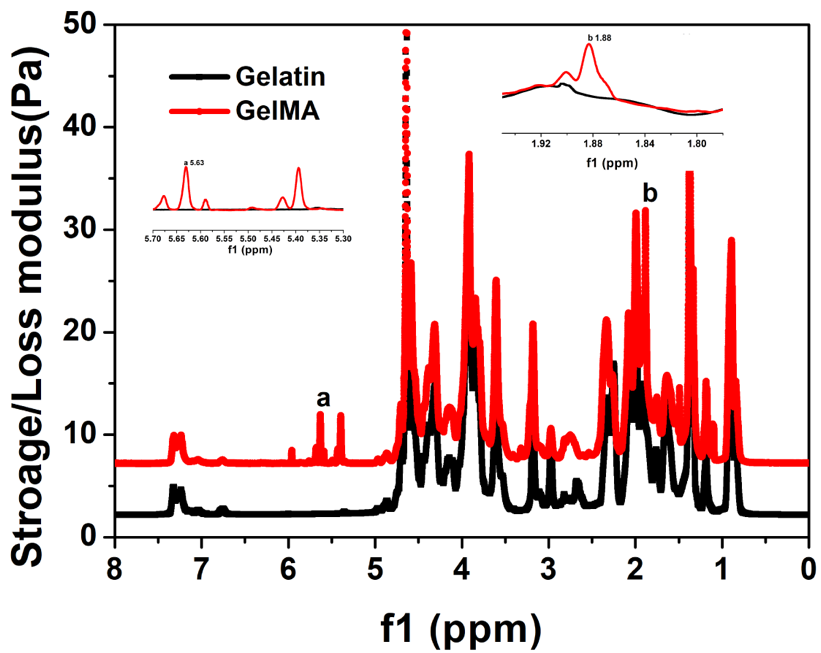

**Figure S2.** The ^1^H NMR spectra showed new signals appear at δ =1.87 ppm, which corresponded to the methyl group of methacrylic acid, and the peaks at δ = 5.4 ppm and δ = 5.6 ppm were the acrylic protons of methacrylic functions in the spectrum of GelMA. The peak at δ = 7.3 ppm represented the aromatic amino acid residues of gelatin.

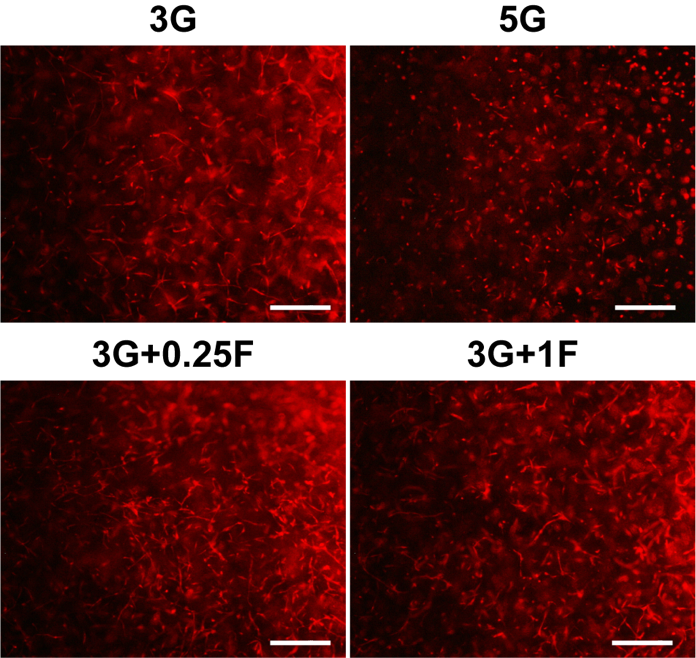

**Figure S3.** RFP-HUVCEs and MSCs encapsulated in corresponding GelMA or GF hydrogels with different percentages showing capillary-like network formation. Scale bar, 100 μm.

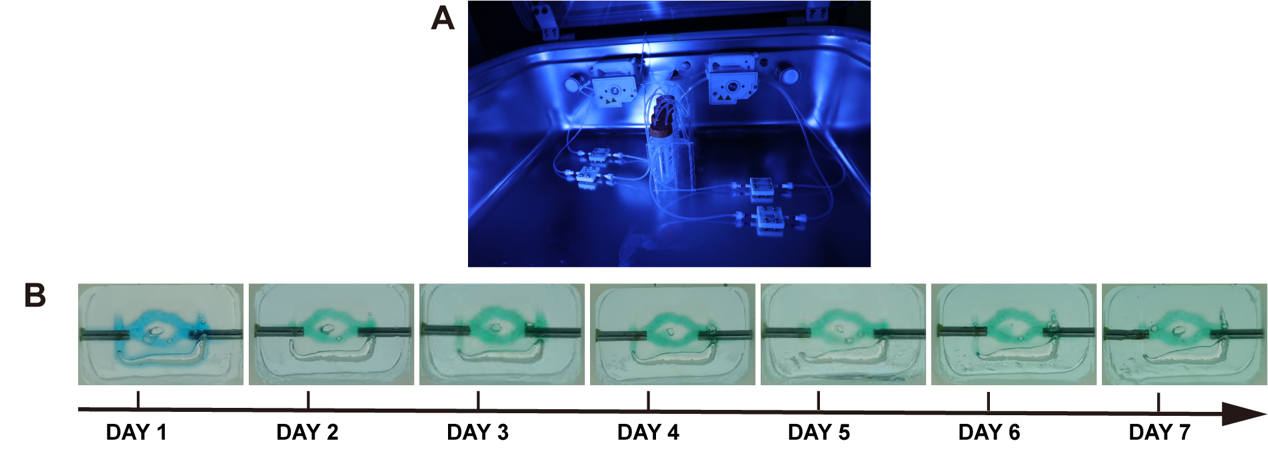

**Figure S4.** A) Images of perfusion equipment showing in-built digital-control peristaltic pump, reservoir of culture medium and several perfusion chambers containing printed tissues. B) “one to two” channel tissue perfused in one weeks. Images showing channel perfused in day 1 to day 7.

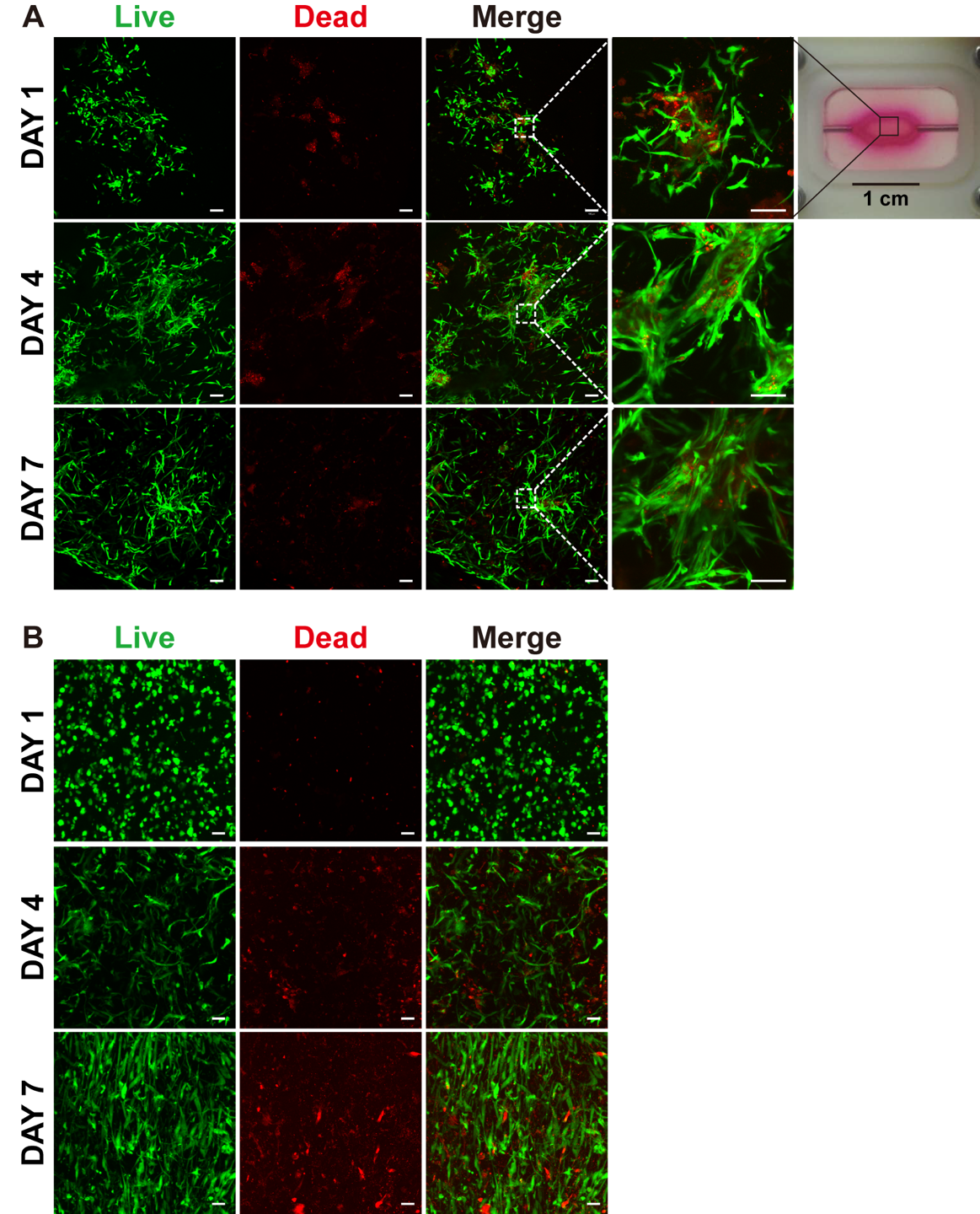

**Figure S5.** Live/dead fluorescence images showing A) HUVECs and MSCs co-cultured aggregates printing. B) HUVECs and MSCs co-printing. Scale bar, 100 μm.

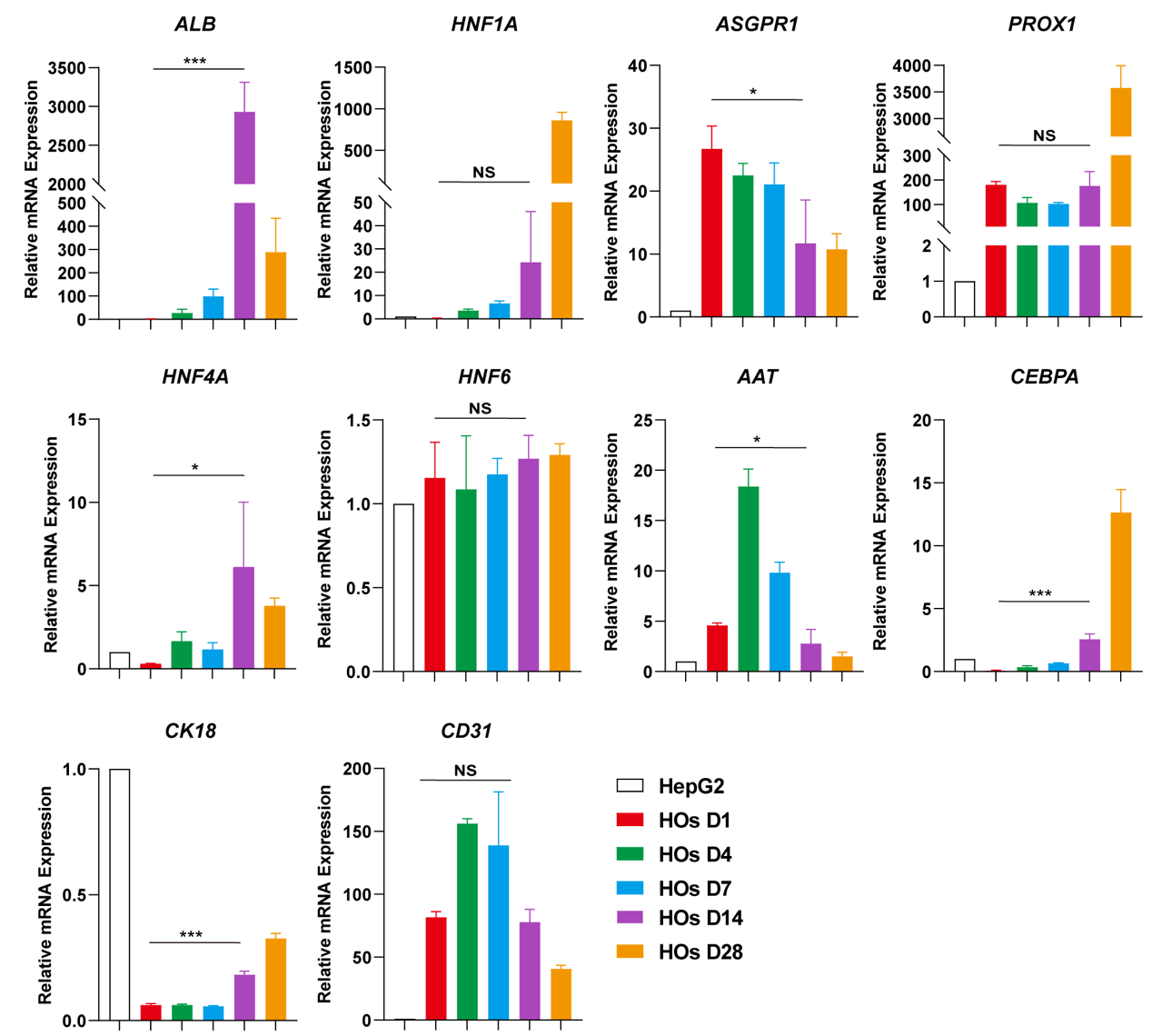

**Figure S6.** Quantitative PCR analysis of the relative hepatic marker gene (*ALB*, *HNF1A*, *ASGPR1*, *PROX1*, *HNF4A*, *HNF6*, *AAT*, *CEBPA*, *CK18*) and the relative vascular marker gene (*CD31*) expression. The data are presented as the mean ± SD. *: p<0.1, **: p<0.01, ***: p<0.001.

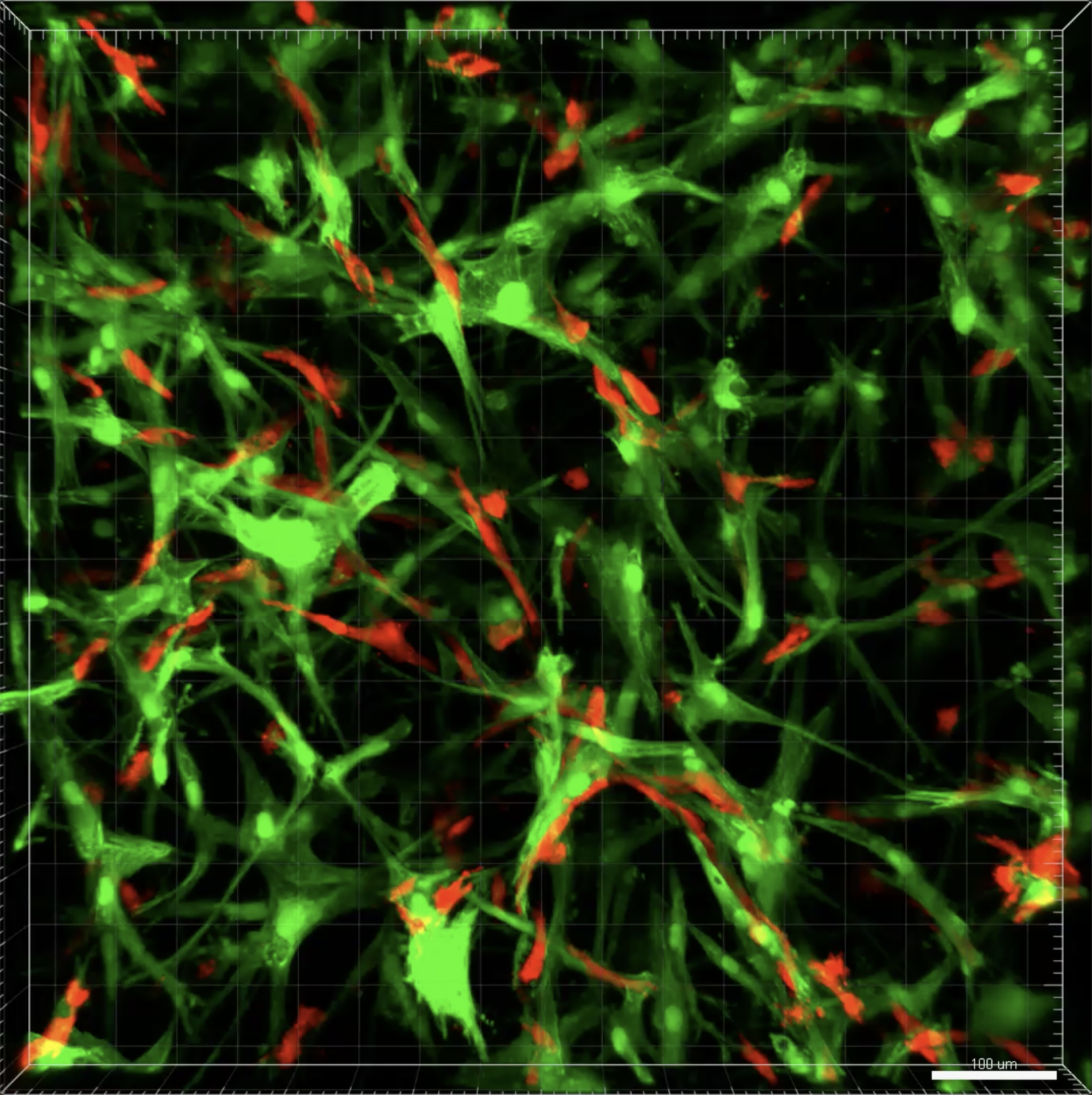

**Movie S1.** 3D reconstruction of constructs from confocal microscopy images showing capillary-like network formation.

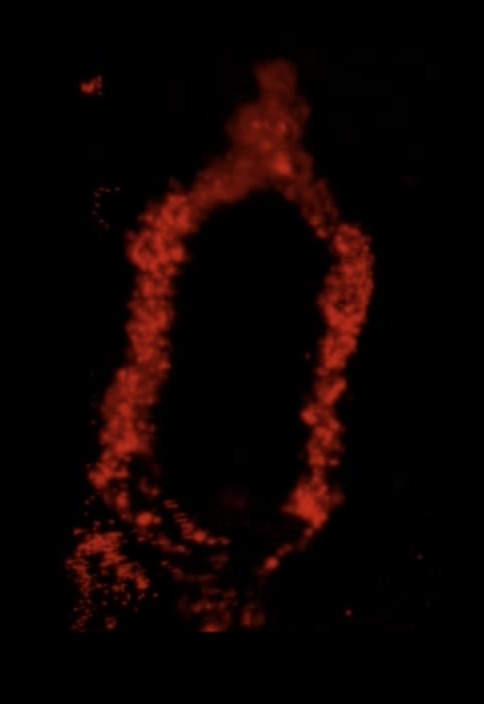

**Movie S2.** Observation of VOs.

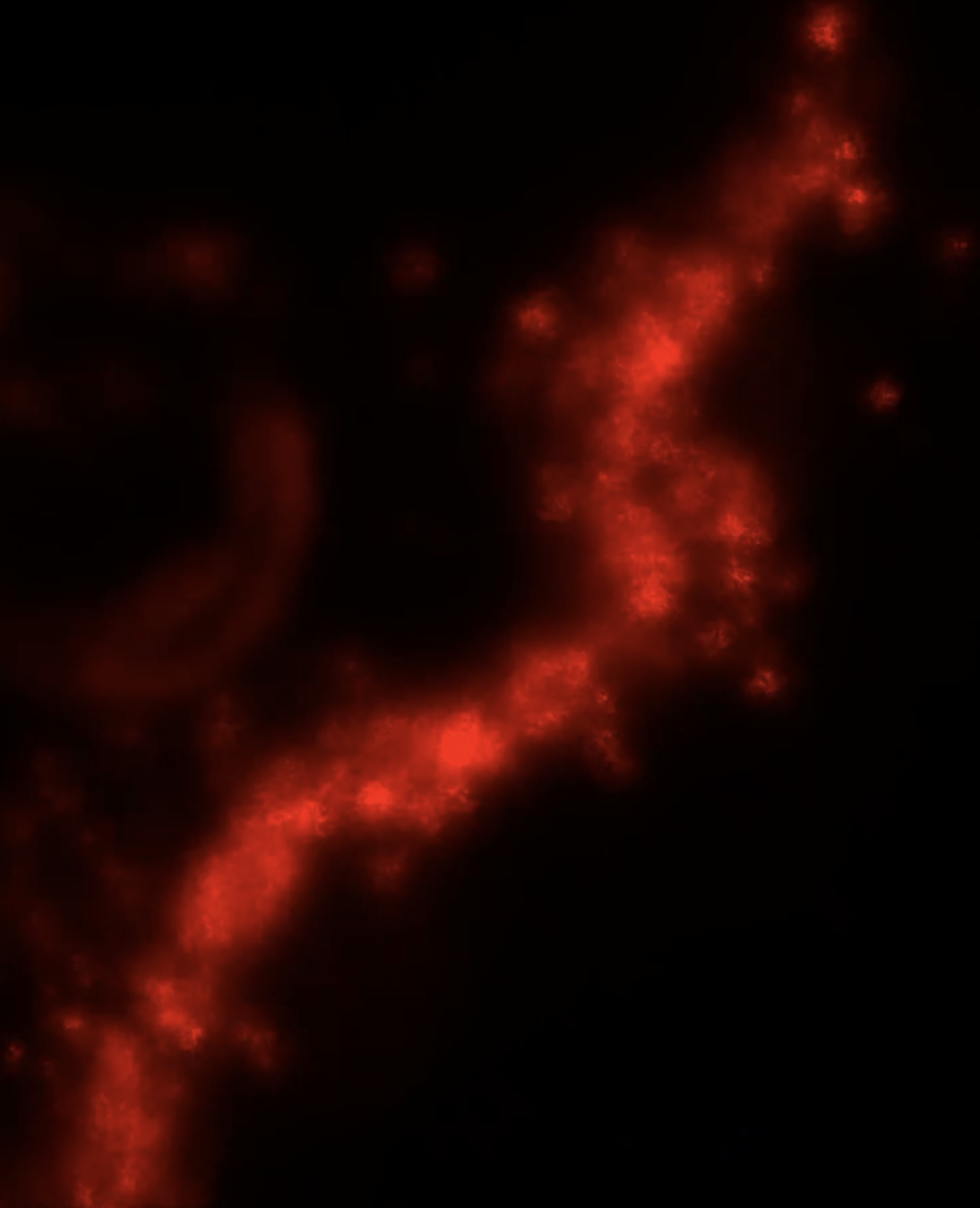

**Movie S3.** VOs perfused with beads.

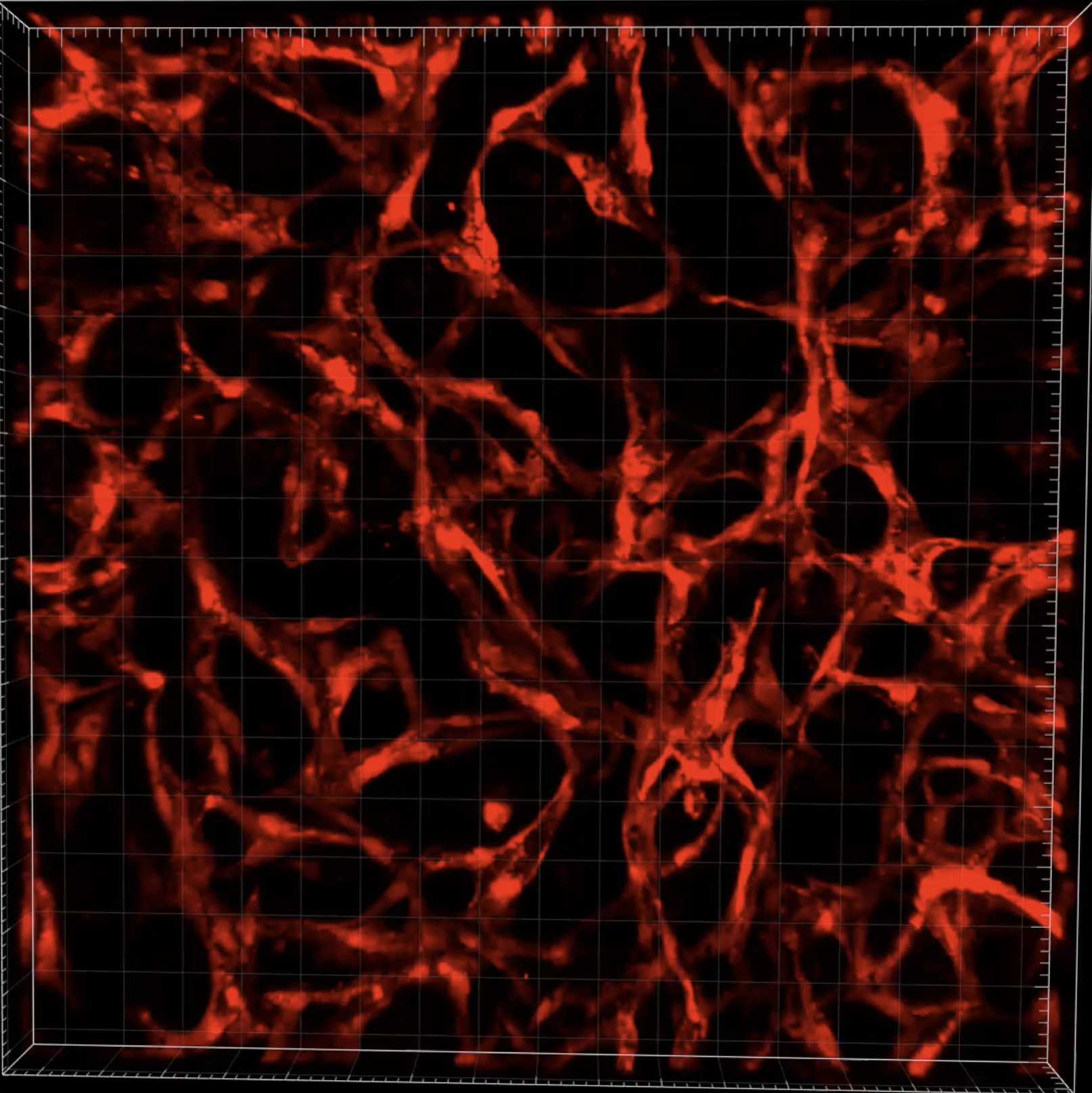

**Movie S4.** Fluorescence composite images displaying vascularized cells in printed GF inks.

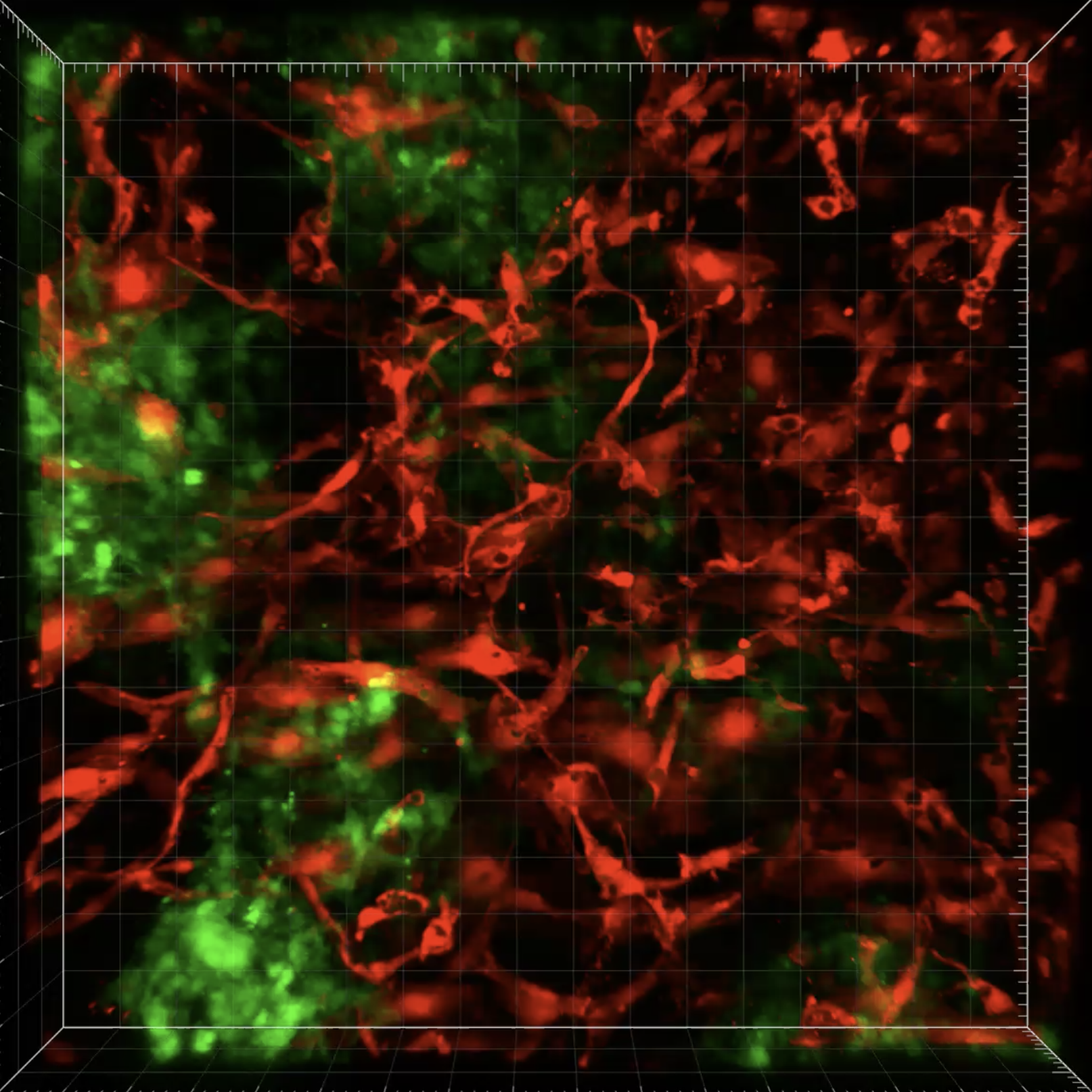

**Movie S5.** Fluorescence images of HOs displaying HAs formed by GFP-HepG2, RFP-HUVECs, and RFP-HUVECs vascularization.

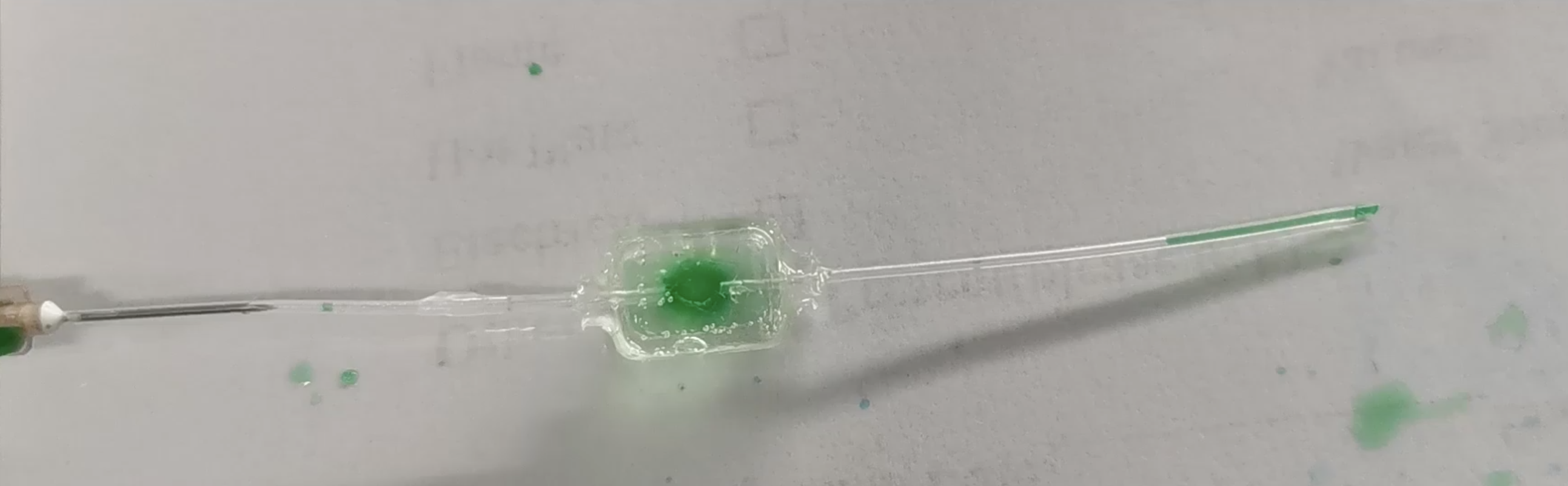

**Movie S6.** “one-to-two” channeled tissues encapsulated in PDMS connected with PU tubes perfused with green paint.

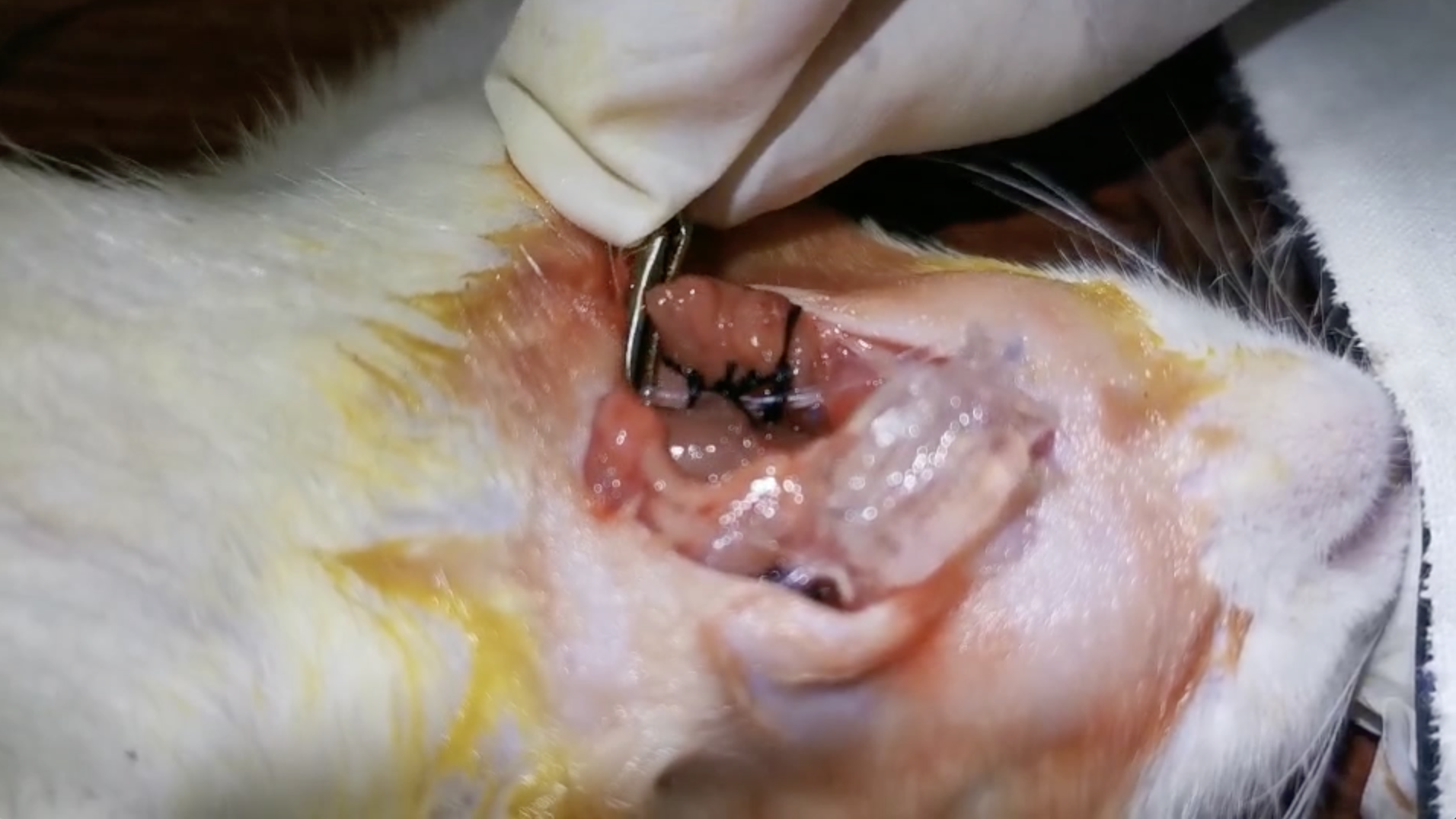

**Movie S7.** Observation of transplanted tissues after establishment of connections with carotid artery and jugular vein showing arterial clip slip and establishment of blood perfusion.

**Table S1.** Primer sequences used in this study.

| Gene name | Forward (5^’^-3’) | Reverse (5’-3’) |
| --- | --- | --- |
| *CD31* | TCTATGACCTCGCCCTCCACAAA | GAACGGTGTCTTCAGGTTGGTATTTCA |
| *ALB* | CCTTTGGCACAATGAAGTGGGTAACC | CAGCAGTCAGCCATTTCACCATAGG |
| *AAT* | TATGATGAAGCGTTTAGGC | CAGTAATGGACAGTTTGGGT |
| *ASGPR1* | GAGAGAGACGTTCAGCAACTTC | GGGACTCTAGCGACTTCATCTT |
| *CK18* | TCGCAAATACTGTGGACAATGC | GCAGTCGTGTGATATTGGTGT |
| *HNF4A* | CACGGGCAAACACTACGGT | TTGACCTTCGAGTGCTGATCC |
| *PROX1* | AAAGGACGGTAGGGACAGCAT | CCTTGGGGATTCATGGCACTAA |
| *HNF6* | CTCTGCTCCTCCTGTTCGAC | CGACCAAATCCGTTGACTCC |
| *HNF1A* | GAACATGGGAAGGATAGAGGCA | GTAGAGTTCGACGCTGGACAT |
| *CEBPA* | TGGACAAGAACAGCAACGAG | TCATTGTCACTGGTCAGCTC |
| *GAPDH* | CTCTGCTCCTCCTGTTCGAC | CGACCAAATCCGTTGACTCC |
